# Defining the RNA Interactome by Total RNA-Associated Protein Purification

**DOI:** 10.1101/436253

**Authors:** Vadim Shchepachev, Stefan Bresson, Christos Spanos, Elisabeth Petfalski, Lutz Fischer, Juri Rappsilber, David Tollervey

## Abstract

UV crosslinking can be used to identify precise RNA targets for individual proteins, transcriptome-wide. We sought to develop a technique to generate reciprocal data, identifying precise sites of RNA-binding proteome-wide. The resulting technique, <u>t</u>otal <u>R</u>NA-<u>a</u>ssociated <u>p</u>rotein <u>p</u>urification (TRAPP), was applied to yeast (*S. cerevisiae*) and bacteria (*E. coli*). In all analyses, SILAC labelling was used to quantify protein recovery in the presence and absence of irradiation. For *S. cerevisiae*, we also compared crosslinking using 254 nm (UVC) irradiation (TRAPP) with 4-thiouracil (4tU) labelling combined with ~350 nm (UVA) irradiation (PAR-TRAPP). Recovery of proteins not anticipated to show RNA-binding activity was substantially higher in TRAPP compared to PAR-TRAPP. As an example of preferential TRAPP-crosslinking, we tested enolase (Eno1) and demonstrated its binding to tRNA loops *in vivo*. We speculate that many protein-RNA interactions have biophysical effects on localization and/or accessibility, by opposing or promoting phase separation for highly abundant protein. Homologous metabolic enzymes showed RNA crosslinking in *S. cerevisiae* and *E. coli*, indicating conservation of this property. TRAPP allows alterations in RNA interactions to be followed and we initially analyzed the effects of weak acid stress. This revealed specific alterations in RNA-protein interactions; for example, during late 60S ribosome subunit maturation. Precise sites of crosslinking at the level of individual amino acids (iTRAPP) were identified in 395 peptides from 155 unique proteins, following phospho-peptide enrichment combined with a bioinformatics pipeline (Xi). TRAPP is quick, simple and scalable, allowing rapid characterization of the RNA-bound proteome in many systems.

## INTRODUCTION

Interactions between RNA and proteins play key roles in many aspects of cell metabolism. However, the identification of protein-RNA interaction sites has long been challenging, particularly in living cells. Individual protein-RNA interactions can be characterized, if known, by mutagenic and biochemical approaches but this has always been labor-intensive. The difficulty is compounded by the fact that many interactions do not fall within characterized interaction domains, and even apparently well-characterized RNA-binding domains can show multiple modes of RNA interaction (Clery et al., 2013), making detailed predictions less reliable. For large-scale characterization of protein-RNA interactions, a significant advance was the development of RNA immunoprecipitation (RIP) with or without formaldehyde crosslinking, allowing the identification of RNAs associated with target proteins, although not the site of association (Gilbert et al., 2004; Gilbert and Svejstrup, 2006; Huang et al., 2005; Hurt et al., 2004; Motamedi et al., 2004; Niranjanakumari et al., 2002). Subsequently, UV crosslinking approaches were developed that allow accurate identification of the binding sites for individual proteins on RNA molecules, transcriptome-wide (Bley et al., 2011; Doneanu et al., 2004; Granneman et al., 2009; Granneman et al., 2010; Maly et al., 1980; Mital et al., 1993; Rhode et al., 2003; Urlaub et al., 2000; Van Nostrand et al., 2016; Wagenmakers et al., 1980). Two major approaches have been adopted for crosslinking. Short wavelength, 254 nm UVC irradiation can directly induce nucleotide-protein crosslinking and was used in initial crosslinking and immunoprecipitation (CLIP) analyses, as well as CRAC analyses in yeast. Subsequently, photoactivatable ribonucleoside-enhanced crosslinking and immunoprecipitation (PAR-CLIP) was developed, in which 4-thiouracil is fed to the cells and incorporated into nascent transcripts, allowing RNA-protein crosslinking to be induced by longer wavelength, ~350-365nm UVA irradiation. Both approaches have been used extensively in many systems.

The reciprocal analyses of proteins that are bound to RNA were more difficult to develop, at least in part because proteomic approaches do not provide the amplification offered by PCR. However, UV crosslinking and RNA enrichment have been used to successfully identify many poly(A)^+^ RNA binding proteins present in human cells and other systems (Baltz et al., 2012; Castello et al., 2012; Kwon et al., 2013). This technique was an important advance but, like RIP, identifies the species involved but not the site of interaction, and is limited to mature mRNAs. To identify the total RNA-bound proteome, the approach of 5-ethynyluridine (EU) labelling of RNAs followed by biotin ligation using the click reaction (RICK) was recently developed (Bao et al., 2018; Huang et al., 2018) as well as approaches based on phase separation, in which RNA-protein conjugates are recovered from an aqueous/phenol interface (Garcia-Moreno et al., 2018; Gatto et al., 2018; Trendel et al., 2018). In addition, MS analyses have been developed to identify the precise amino-acid at the site of RNA-protein crosslinking (Kramer et al., 2014).

The TRAPP techniques for identification of RNA-protein interaction are simple, cost-effective, quantitative and readily scalable. In addition, they permit identification of the peptide, and indeed the amino acid, that is crosslinked to RNA during UV irradiation in living cells. We applied TRAPP to identify RNA-binding proteins from yeast and *E. coli* following crosslinking in actively growing cells. These approaches will allow reliable and accurate characterization of the steady state protein-RNA interactome as well as it’s dynamics upon stress exposure in many systems.

## RESULTS

### Development of the TRAPP protocol

In all TRAPP techniques, we initially covalently linked all RNAs to associated proteins by UV crosslinking in actively growing cells. Detailed protocols for each of the TRAPP workflows are given in Materials and Methods.

In the TRAPP approach, ~750ml cultures of actively growing yeast were irradiated at 254 nm (UVC) using the workflow outlined in Figure 1A. To quantify protein recovery by mass spectrometry in the presence and absence of UV crosslinking, the analyses incorporated <u>s</u>table isotope labelling with <u>a</u>mino acids in <u>c</u>ell culture (SILAC) (Ong et al., 2002) combined with the Maxquant software package (Cox and Mann, 2008). For this, yeast strains that were auxotrophic for lysine and arginine (*lys9*Δ*0*, *arg4*Δ*0*) were grown in the presence of lysine and arginine or [^13^C_6_]-lysine plus [^13^C_6_]-arginine. Label swaps between crosslinked and non-crosslinked samples were used to confirm that the labelling did not affect the outcome of the analyses (Figures S1A-S1C). Labeled and unlabeled cells were mixed prior to cell lysis.

**Figure 1.**
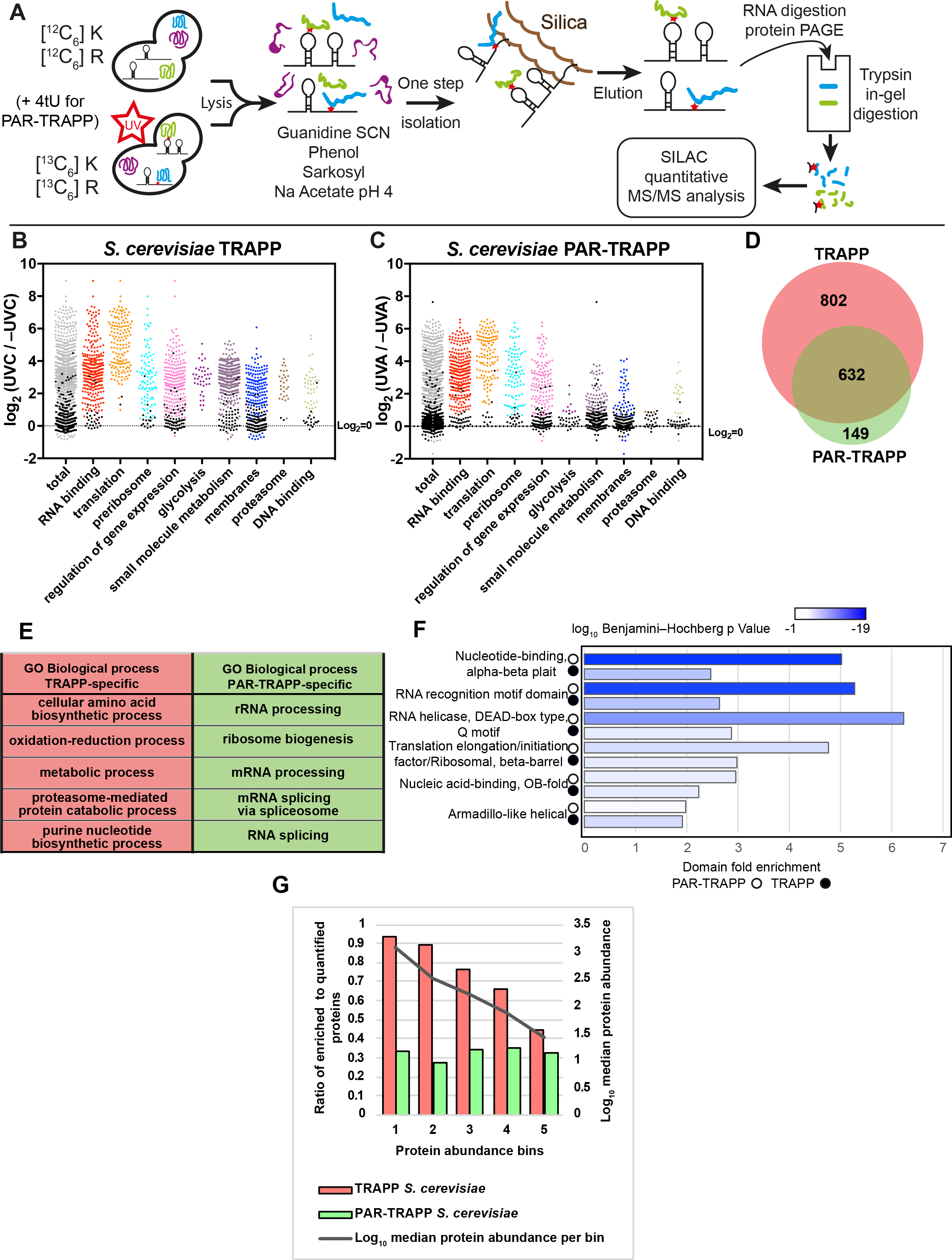
TRAPP and PAR-TRAPP reveal the yeast RBPome. (A) TRAPP and PAR-TRAPP workflows used to identify RNA-interacting proteins with SILAC MS-MS. See the main text for details. (B) Scatter plot of Log_2_ SILAC ratios +UVC/−UVC for *S. cerevisiae* proteins, quantified with TRAPP. Proteins were subdivided based on the indicated GO term categories. Proteins, belonging to GO terms, "membrane" and "DNA binding" do not contain proteins mapping to GO terms "RNA metabolic process", "RNA binding", "ribonucleoprotein complex". Black dots represent proteins that failed to pass statistical significance cut-off (P value adjusted <0.05). (C) Scatter plot of Log_2_ SILAC ratios +UVA/−UVA for *S. cerevisiae* proteins, quantified with PAR-TRAPP. Proteins were subdivided based on the indicated GO term categories. Proteins, belonging to GO terms "small molecule metabolism", "membrane" and "DNA binding" do not contain proteins mapping to GO terms "RNA metabolic process", "RNA binding", "ribonucleoprotein complex". Black dots represent proteins that failed to pass statistical significance cut-off (P value adjusted <0.05). (D) Venn diagram showing the overlap between proteins identified in TRAPP and PAR-TRAPP. (E) 5 most enriched GO terms amongst proteins identified only in TRAPP or exclusively in PAR-TRAPP. (F) 6 most significantly enriched domains (lowest P value) in PAR-TRAPP-identified proteins were selected if the same domain was enriched amongst TRAPP-identified proteins. Domain fold enrichment in the recovered proteins is plotted on the x axis, while colour indicates log_10_ Benjamini–Hochberg adjusted P value. (G) Proteins quantified in both TRAPP and PAR-TRAPP were filtered to remove proteins annotated with GO terms "RNA metabolic process", "RNA binding", "ribonucleoprotein complex". The remaining proteins were split into 5 bins by abundance (see materials and methods). For each bin the ratio between enriched to total quantified proteins was calculated as well as median protein abundance as reported by PaxDb.

Briefly, irradiated cells were harvested by centrifugation, resuspended in buffer containing 2M guanidine thiocyanate plus 50% phenol, and lysed by beating with zirconia beads. The lysate was cleared by centrifugation and adjusted to pH4 by adding 3M sodium acetate. The cleared lysate was incubated with silica beads in batch for 60min. The beads were extensively washed in denaturing buffer; first in buffer containing 4M guanidine thiocyanate, 1M sodium acetate pH4.0, then in low salt buffer with 80% ethanol. Nucleic acids and bound proteins were eluted from the column in Tris buffer. The eluate was treated with RNase A + T1 to degrade RNA and proteins were resolved in a polyacrylamide gel in order to remove degraded RNA, followed by in-gel trypsin digestion and LC-MS/MS analysis.

The PAR-TRAPP approach is similar to TRAPP, except that cells were metabolically labeled by addition of 4-thiouracil (4tU) to the culture (final concentration 0.5mM) for 3h prior to irradiation, in addition to SILAC labelling. 4tU is rapidly incorporated into RNA as 4-thiouridine, thus sensitizing RNA to UVA crosslinking. However, 4tU strongly absorbs UV irradiation and confers considerable UV-resistance on the culture (Figure S1D). Cells were therefore rapidly harvested by filtration and resuspended in medium lacking 4tU immediately prior to irradiation at ~350 nm (UVA). Previous analyses using labelling with 4tU and UVA irradiation in yeast and other systems have generally involved significant crosslinking times (typically 30min in a Stratalinker with 360nm UV at 4mJ sec^-1^ cm^-2^), raising concerns about changes in RNA-protein interactions during this extended period of irradiation. We therefore constructed a crosslinking device (Figure S1E) that delivers a substantially increased UV dose, allowing irradiation times of only 38 sec to be used to deliver the equivalent dose of 7.3J cm^-2^. Subsequent treatment was as described above for TRAPP.

In principal, silica binding can enrich both RNA and DNA bound proteins, although UV crosslinking to dsDNA is expected to be inefficient (Angelov et al., 1988). To compare recovery of proteins bound to RNA versus DNA, samples recovered following initial silica binding and elution were treated with either DNase I, RNase A + T1 or cyanase (to degrade both RNA and DNA) and then rebound to silica, as outlined in Figure S2A. Following washing and elution from the silica column, proteins were separated by SDS-polyacrylamide gel electrophoresis and visualized by silver staining (Figures S2B-S2C). Nucleic acids were separated by agarose gel electrophoresis and visualized by SYBR safe staining (Figure S2D) to confirm degradation. Degradation of DNA had little effect on protein recovery, whereas this was substantially reduced by RNase treatment. The predominant proteins correspond to the added RNases. We conclude that the TRAPP protocol predominately recovers RNA-bound proteins. However, even following treatment with cyanase or in the absence of UV irradiation (-UVC), some proteins were recovered following RNA binding to silica. These proteins apparently bind RNA in the absence of crosslinking, even following denaturation, likely due to the mixed mode of binding, involving both hydrophobic interactions with nucleotide bases and charge interactions with the phosphate backbone. These will be under-estimated in SILAC quantitation of TRAPP analyses, potentially generating a small number of false negative results.

Maxquant quantitation initially failed to return a value for many peptides in TRAPP MS/MS data, predominately due to the absence of a detectable peptide in the −UV samples for the SILAC pairs (Figure S3A-F). These “missing” peptides were strongly enriched for known RNA binding proteins, presumably because low abundance proteins that are efficiently purified are detected following crosslinking but not in the negative control. We addressed this problem by imputation of the missing values using the imputeLCMD R package (see Materials & Methods and Figures S3G-S3O) (Lazar et al., 2016).

TRAPP in *S. cerevisiae* identified 1,434 proteins significantly enriched by UVC crosslinking, of which 1,360 were enriched more than 2-fold (Table S1). Proteins annotated with the GO term “translation” had the highest average fold enrichment, which is expected in the total RNA-interacting proteome (Figure 1B). Several reports have recently proposed that proteins involved in cellular metabolism can interact with RNA. Consistent with this we observed significant enrichment for proteins annotated with GO term "small molecule metabolic process" (Figure 1B). Furthermore, 341 proteins enriched in TRAPP were annotated in the Yeast Metabolome Database (YMDB) (Ramirez-Gaona et al., 2017) to be enzymes or transporters associated with pathways of intermediary metabolism (table S1). In particular, the majority of enzymes involved in glycolysis and/or gluconeogenesis were identified in TRAPP (Figure S4) with average enrichment of more than 4-fold. In addition, most of the structural components of the proteasome and many membrane-associated proteins were identified as RNA-binders by TRAPP.

Applying PAR-TRAPP identified 2-fold fewer significantly enriched proteins than found with TRAPP (Table S2). However, PAR-TRAPP recovered a notably higher proportion of characterized RNA-binding proteins relative to proteins with less obvious connection to RNA. Only 81 proteins implicated in intermediary metabolism by the YMDB were identified among the PAR-TRAPP hits (relative to 341 in TRAPP) (Table S2). Furthermore, proteins annotated with the GO terms "glycolysis", "membrane part" and "proteasome" were substantially reduced relative to TRAPP (Figure 1C).

We performed GO term analyses on proteins that were exclusively found enriched in TRAPP or PAR-TRAPP (Figure 1D-E). The most enriched GO terms for TRAPP-specific proteins were related to metabolic processes and the proteasome, whereas PAR-TRAPP-specific proteins featured rRNA processing, mRNA processing and RNA splicing. Furthermore, while both TRAPP and PAR-TRAPP demonstrated over-representation for known RNA-interacting domains within the enriched proteins (Table S3), in PAR-TRAPP this trend was more pronounced both in terms of higher fold enrichment and in terms of enrichment p-values (Figure 1F).

We assessed the correlation between cellular abundance and the likelihood of being reported as RNA interacting protein by TRAPP and PAR-TRAPP. Strikingly almost 90% of proteins within the two most abundant bins were scored as enriched in +UV by TRAPP, with a gradual decline of the number amongst less abundant proteins, whereas no such correlation was observed in PAR-TRAPP data (Figure 1G). Only confidently identified proteins were included in each analysis, so this is unlikely to reflect a detection bias. Since the trend is not observed in PAR-TRAP, it also seems unlikely to reflect a systematic bias towards higher confidence (lower p values) for abundant proteins. This suggests that abundant proteins have features that predispose them towards RNA association. As discussed below (see Discussion), TRAPP is predicted to better recover transient interactions than PAR-TRAPP, due to differences in the UV-nucleotide interactions involved. We therefore postulate that highly abundant proteins have systematically acquired features that favor transient RNA interaction.

We further compared the RNA interactors identified in PAR-TRAPP with the published results of poly(A) capture (Beckmann et al., 2015), considering only proteins identified with 7.2 J cm^-2^ UVA plus 4tU, as in PAR-TRAPP. There was a substantial overlap between proteins reported by PAR-TRAPP and poly(A) capture techniques and this was increased by including the results of TRAPP (Figures 2A). First, we compared protein fold enrichment for selected GO terms in TRAPP versus poly(A) capture (Figures 2B-2C). For proteins included under the terms "RNA binding", "translation", "preribosome" and "mRNA binding" TRAPP had a higher dynamic range compared to poly(A) capture (Figure 2B). This was notable for "mRNA binding" as poly(A) capture was expected to perform particularly well for these proteins. In contrast, the GO terms "membrane", "glycolysis" and "small molecule metabolic processes" showed higher average enrichment in poly(A) capture than PAR-TRAPP. This raises the possibility that at least some RNA-interacting proteins of intermediary metabolism interact with mRNA species.

**Figure 2.**
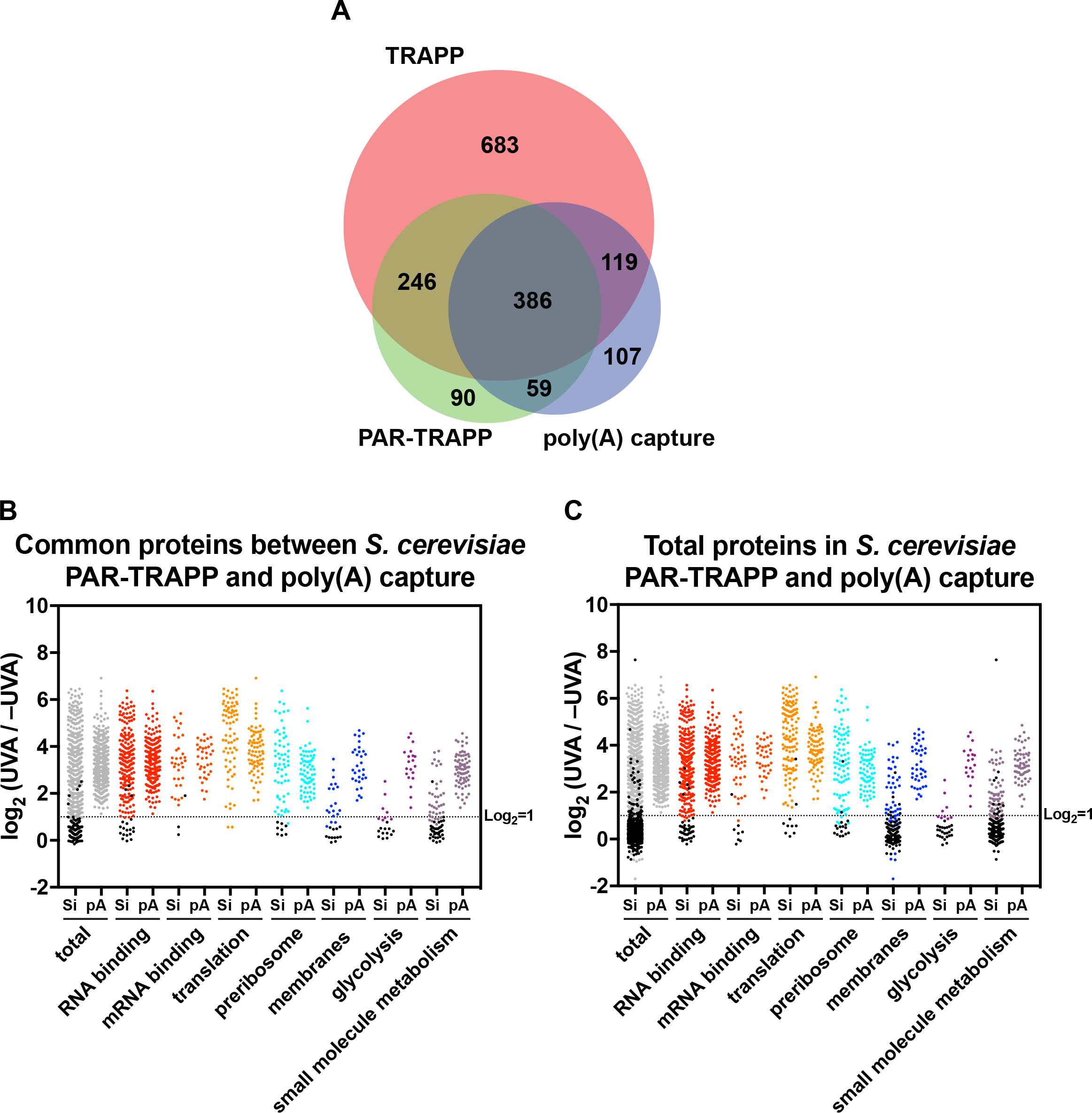
The yeast total RBPome compared to poly(A) RNA RBPome. A: Venn diagram showing the overlap between proteins identified in PAR-TRAPP, poly(A) capture and TRAPP. (B) Scatter plot of Log_2_ SILAC ratios +UVA/−UVA for *S. cerevisiae* proteins common between PAR-TRAPP and Log_2_ +UVA/−UVA fold enrichment for poly(A) capture technique. Proteins were subdivided based on the indicated GO term categories. Proteins, belonging to GO terms, "membrane" and "small molecule metabolism" do not contain proteins mapping to GO terms "RNA metabolic process", "RNA binding", "ribonucleoprotein complex". Black dots represent proteins that failed to pass statistical significance cut-off (P value adjusted <0.05). (C) Scatter plot of Log_2_ SILAC ratios +UVA/−UVA for all *S. cerevisiae* proteins identified in PAR-TRAPP and Log_2_ +UVA/−UVA fold enrichment for poly(A) capture technique. Labelling is as in panel B.

### TRAPP identification of RNA-binding proteins in gram-negative bacteria

Sequence independent binding to silica, combined with aggressive, highly denaturing lysis, makes TRAPP potentially applicable to any organism. To confirm this, we applied TRAPP to *E. coli*, which is not amenable to proteome capture through poly(A) RNA isolation. We used TRAPP, as we could not identify a 4tU concentration that allowed both growth and efficient RNA-protein crosslinking. TRAPP in *E. coli* identified 1,106 significantly enriched proteins, of which 1,089 were enriched more than 2-fold (Table S4). Enrichment by GO term categories was similar to yeast TRAPP data (Figure 3A), with proteins involved in translation showing the highest average enrichment compared to −UV. Other major RNA-binding proteins showed enrichment similar to ribosomal proteins, including the RNA chaperone Hfq and the cold-shock proteins CspC, CspE, CspA and CspD.

**Figure 3.**
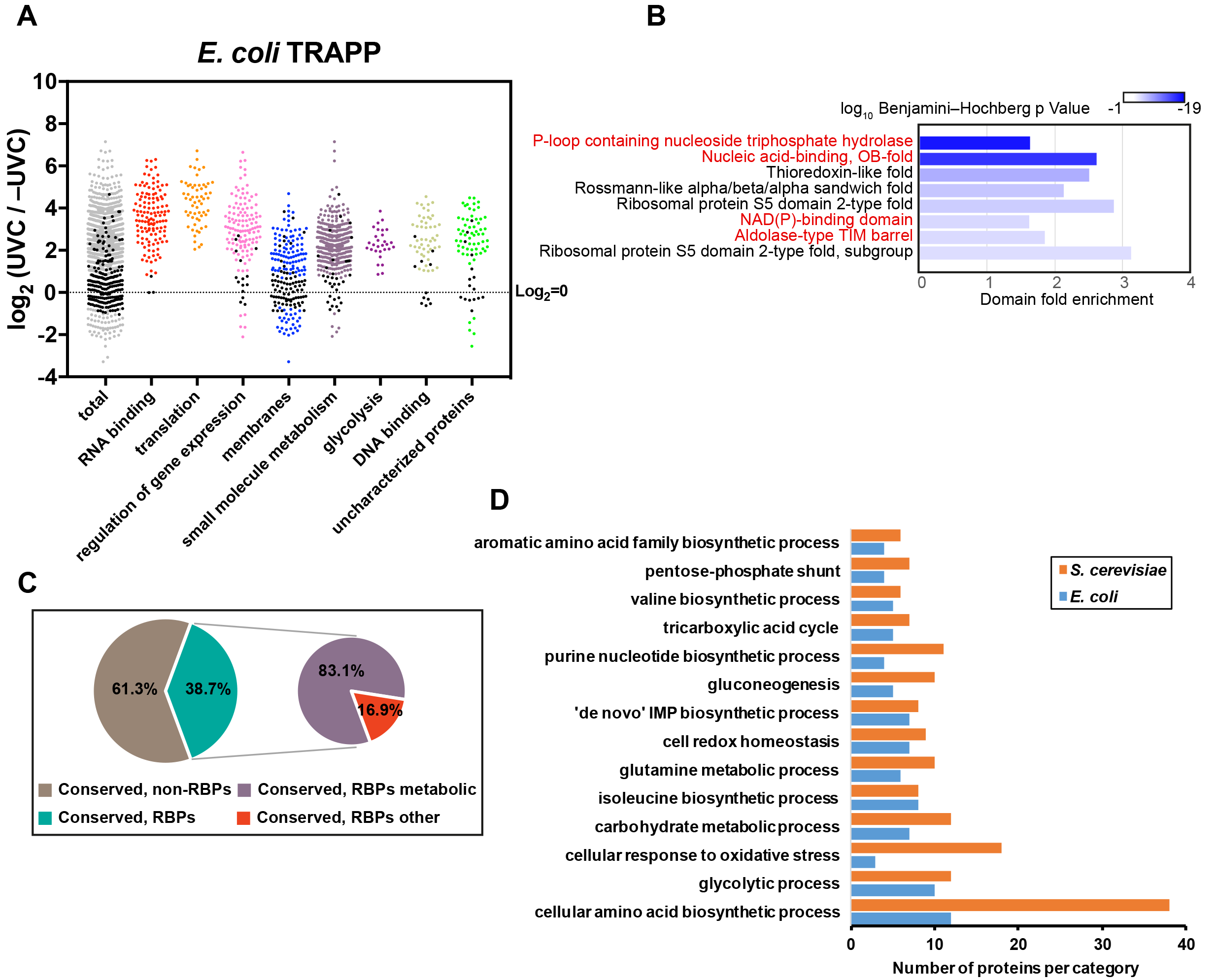
TRAPP reveals RNA binding proteins conserved from *E. coli* to *S. cerevisiae*. (A) Scatter plot of Log_2_ SILAC ratios +UVC/−UVC for *E. coli* proteins, quantified with TRAPP. Proteins were subdivided based on the indicated GO term categories. Proteins, belonging to GO terms, "membrane", "small molecule metabolism" and "DNA binding" do not contain proteins mapping to GO terms "RNA metabolic process", "RNA binding", "ribonucleoprotein complex". Black dots represent proteins that failed to pass statistical significance cut-off (P value adjusted <0.05). Uncharacterized proteins are proteins annotated as " Putative uncharacterized protein" in the Uniprot database. (B) Most significantly enriched protein domains (lowest P value) in *E. coli* TRAPP-identified proteins. Fold enrichment of the indicated domain amongst the recovered proteins is plotted on the x axis, while color indicates log_10_ Benjamini–Hochberg adjusted P value. Domains found enriched in the yeast TRAPP data are labelled with red colour. (C) Pie chart of Inparanoid 8.0 database orthologous clusters between *S. cerevisiae* and *E. coli*. For a cluster to be labelled as conserved RNA interacting ("conserved, RBPs"), it was required to contain at least one bacterial and one yeast protein enriched in TRAPP. "Conserved, RBPs metabolic" are clusters where at least one protein in yeast or bacteria is identified in the YMDB or in ECMDB databases respectively (see materials and methods). (D) Results of GO term enrichment analysis for conserved RNA interacting proteins of intermediary metabolism from yeast and bacteria, identified in panel C.

A number of DNA-binding proteins, including Fis, HupA and HupB (HU), H-NS, Dps, StpA, IhfA and IhfB, showed enrichment of more than 11-fold. However, several of these proteins were reported to also interact with RNA; Hu (Macvanin et al., 2012), H-NS (Brescia et al., 2004), Dps (Windbichler et al., 2008) and StpA (Zhang et al., 1995). In addition, Helix-Turn-Helix transcription factors were significantly enriched, as were components of the KEGG pathway "nucleotide excision repair". We cannot exclude the possibility that TRAPP in *E. coli* detected a subset of DNA binding proteins.

Domain enrichment analysis (Figure 3B) showed that *E. coli* proteins identified by TRAPP were enriched in known RNA-binding domains. However, domains not clearly related to RNA were also strongly enriched, notably NAD(P)-binding domains and the Aldolase-type TIM barrel domain which were also enriched amongst the TRAPP proteins in yeast. Consistent with this, GO analyses identified many significantly enriched proteins categorized as "small molecule metabolism" (Figure 3A). Thioredoxin-like fold domains were enriched (Figure 4B), as previously found in human cells (Castello et al., 2016), suggesting that non-canonical protein-RNA interactions might be highly conserved. However, we observed a similar correlation between abundance and TRAPP enrichment in *E. coli* data (data not shown).

**Figure 4.**
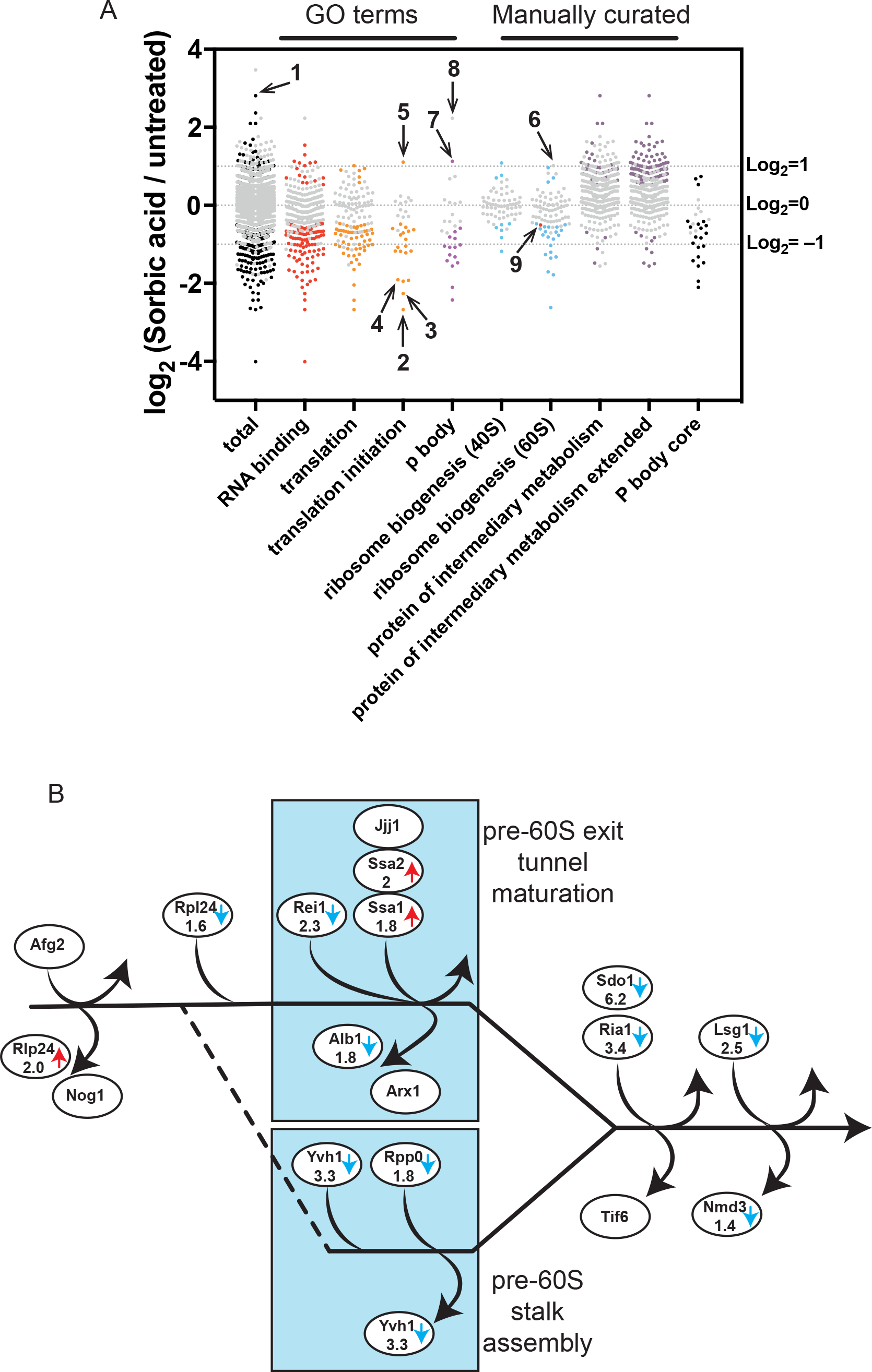
TRAPP reveals the dynamics of RBPome upon stress. (A) Scatter plot of Log_2_ SILAC ratios +Sorbic/−Sorbic for S. cerevisiae proteins, quantified with TRAPP. Grey points represent proteins showing no statistically significant change upon sorbic acid exposure in TRAPP, while proteins changing significantly (P value adjusted <0.05) are labelled with other colors. Only proteins observed as RNA interacting in PAR-TRAPP were included in the analysis, except for proteins in the category "protein of intermediary metabolism extended", for which this criterion was dropped. Proteins annotated with GO term categories "RNA binding", "translation", "translation initiation", "p body" and "small molecule metabolism" are displayed together with proteins annotated in literature-curated lists: "ribosome biogenesis 40S", "ribosome biogenesis 60S" (Woolford and Baserga, 2013). Proteins, belonging to categories "protein of intermediary metabolism" and "protein of intermediary metabolism extended" are yeast enzymes and transporters of intermediary metabolism, obtained from YMDB and further filtered to remove aminoacyl-tRNA synthetases. "P body core" category contains proteins identified as core components of p bodies in yeast (Buchan et al., 2010). Numbers label the following protein on the chart: 1 - Rtc3; 2 – Tif3; 3 – Rpg1; 4 – Tif35; 5 – Gcd11; 6 – Rlp24; 7 – Ssd1; 8 – Rbp7; 9 – Nmd3. (B) The cytoplasmic phase of large subunit maturation in yeast (adopted from (Lo et al., 2010)). Proteins altered in abundance in PAR-TRAPP data upon sorbic acid exposure are indicated with arrows. Blue arrow denotes decrease, while red arrows indicate increase in PAR-TRAPP recovery upon stress. For proteins passing the statistical significance cut-off (P value adjusted <0.05), fold change is indicated.

This observation prompted us to analyze orthologous proteins between *S. cerevisiae* and *E. coli*. The Inparanoid 8 database (Sonnhammer and Östlund, 2015) reports 460 orthologous clusters between the two species, 38.7% of which had at least one bacterial and one yeast protein enriched in TRAPP (Figure 3C). Most of these clusters contained metabolic enzymes annotated in the corresponding metabolome databases for budding yeast and *E. coli*, so we extracted all of the conserved TRAPP proteins and performed a KEGG pathway enrichment analysis. This identified a number of enriched pathways including “glycolysis and gluconeogenesis”a (Figure S5) and “purine biosynthesis” (Figure S6), as well as “pentose phosphate” and multiple amino acid biosynthetic pathways (data not shown).

### TRAPP reveals dynamic changes in RNA-protein interaction following stress

Cellular RNPs can be dramatically remodeled under altered metabolic conditions. For the initial analyses we applied weak acid stress, which results in rapid cytoplasm acidification, as a drop in intracellular pH was reported to be a common feature of response to a variety of stresses (Weigert et al., 2009). Furthermore, reduced intracellular pH directly contributes to phase separation of poly(A) binding protein Pab1 into gel-like structures (Riback et al., 2017). We applied PAR-TRAPP as it gave a greater dynamic range of RNA-protein interactions following stress, relative to TRAPP (data not shown). SILAC-labelled yeast cells were treated with 6 mM sorbic acid for 16 min, to induce weak acid stress, followed by PAR-TRAPP. Only proteins previously identified as RNA-interacting by PAR-TRAPP were considered for the initial analysis.

Recovery of the majority of quantified proteins was unchanged upon this brief exposure to sorbic acid (Figure 4A), increasing the likelihood that those alterations observed reflect changes in RNA binding rather than protein expression. Statistically significant changes of >2-fold were observed for 123 proteins; 100 showed reduced RNA association, while 23 were increased (Table S6). Analysis of translation related proteins indicted the specific inhibition of translation initiation, with six initiation factors showing >2-fold reduced RNA-binding. The most reduced were eIF4B (Tif3) (6.4-fold), eIF3A (Rpg1) (4.8-fold) and eIF3G (Tif35) (3.7-fold). Tif3 is of particular interest in the context of weak acid stress, since *tif3Δ* cells are hyper-sensitive to acetic and propionic acids (Kawahata et al., 2006; Mira et al., 2008) and a subset of yeast mRNAs are specifically sensitive to loss of Tif3 (Sen et al., 2016). We speculate that Tif3 may orchestrate responses to acid stress at the level of translation initiation.

The only translation initiation factor to show increased RNA-binding was the GTPase eIF2G (Gcd11), while the two other subunits of the eIF2 complex, eIF2A (Sui2) and eIF2B (Sui3), were unchanged. The role of eIF2 is to escort initiator methionine tRNA to 40S subunits during translation initiation. Similar enrichment upon sorbic acid exposure was observed for the translation elongation factor and GTPase, eEF1A (Tef1), which escorts tRNAs to elongating ribosomes. However, the results may reflect alternative functions for these GTPases during stress response, involving different interactions with RNA. For example, eEF1A was implicated in tRNA export from the nucleus (Grosshans et al., 2000) and in the inhibition of Tor1 kinase activity, presumably acting together with uncharged tRNAs (Kamada, 2017).

A poorly characterized protein, Rtc3/Hgi1 showed 7-fold increased RNA-binding upon sorbic acid treatment. Rtc3 was previously suggested to enhance translation of stress-response proteins (Gomar-Alba et al., 2012) and Rtc3 over-expression confers resistance to weak acids (Hasunuma et al., 2016).

Ribosome biogenesis also showed apparently specific changes following sorbic acid treatment (Figure 4A). Thirteen ribosome synthesis factors that are loaded onto cytoplasmic pre-60S particles following nuclear export showed reduced RNA interactions in sorbic acid treated cells (Figure 4B). The nuclear-cytoplasmic shuttling, pre-60S factors Rlp24 and Nog1 are the first proteins to be released following nuclear export. RNA binding by Rlp24 was significantly increased in sorbic acid treated cells (numbered 6 in Figure 4A); increased binding was also found for Nog1 but was not statistically significant. Rlp24 binds pre-60S particles in the nucleus but following nuclear export it is exchanged for the ribosomal protein Rpl24, which showed reduced RNA binding. This is mediated by the AAA-ATPase Afg2 (also known as Drg1) (Kappel et al., 2012). Displacement of Rlp24 is proposed to be a key initiation step for cytoplasmic pre-60S maturation (Pertschy et al., 2007). The only other cytoplasmic pre-60S factors, binding after Rlp24 release, with increased RNA binding were the homologous chaperones Ssa1 and Ssa2, potentially reflecting structural abnormalities, while all other quantified pre-60S maturation factors had reduced binding. We propose that weak acid stress induces a specific defect in early, cytoplasmic pre-60S ribosomal subunit maturation (Figure 4B).

Following stress, mRNAs were reported to accumulate in storage and processing structures, termed P-bodies and stress granules (Decker and Parker, 2012), although effects of acid stress were not specifically assessed. It was, therefore, unexpected that reduced RNA interactions were seen for eleven proteins classed under the GO term "P-body". Exceptions included the global translation repressor Ssd1 (Hu et al., 2018; Kurischko et al., 2011), which showed >2-fold increased RNA interaction, and Rpb7, which was identified in PAR-TRAPP exclusively under stress conditions. Rpb7 is a component of RNAPII, but dissociates following stress and participates in mRNA decay (Harel-Sharvit et al., 2010; Lotan et al., 2007), presumably explaining its detection only following sorbic acid treatment.

Amongst the 23 proteins showing > 2-fold increased RNA association following sorbic acid, eleven were enzymes participating in intermediary metabolism annotated in the YMDB (Ramirez-Gaona et al., 2017). In contrast, only two such enzymes were identified among the 100 proteins showing >2-fold decreased RNA binding. Our initial analysis only considered proteins that had been identified by PAR-TRAPP under non-stress conditions. Any proteins that interact with RNA only following exposure to stress would therefore be excluded, as found for Rpb7. To avoid this, we generated an extended list of all significant sorbic acid PAR-TRAPP hits. The extended list included 66 proteins with RNA interactions that increased >2-fold following sorbic treatment, of which 36 were YMDB annotated metabolic enzymes, and 113 proteins with >2-fold reduction, five of which were metabolic enzymes (Table S6 extended list; Figure 4A). We conclude that multiple proteins of intermediary metabolism show substantially increased RNA association upon sorbic acid exposure. Notably, a set of metabolic enzymes were previously reported to accumulate in P-bodies and stress granules during stress response (Cary et al., 2015; Jain et al., 2016). Seven of these enzymes showed increased RNA interactions in PAR-TRAPP following sorbic acid treatment, while none showed reduced interactions.

### Identification of precise RNA-binding sites in proteins by iTRAPP

Titanium dioxide (TiO_2_) enrichment can be used to select RNA-crosslinked peptides species due to the presence of phosphate groups on RNA (Richter et al., 2009). For iTRAPP, cells were incubated with 4tU and irradiated with 350 nm UV, as outlined in Figure 5A. Protein-nucleic acid conjugates were initially purified on silica as in TRAPP. Following elution, RNA was digested using nuclease P1, to leave nucleotide 5′ monophosphate groups, and proteins were fragmented with trypsin. Peptides crosslinked to phospho-nucleotides were recovered by enrichment on Titanium dioxide (TiO_2_) columns and the site of crosslinking was identified by tandem MS.

**Figure 5.**
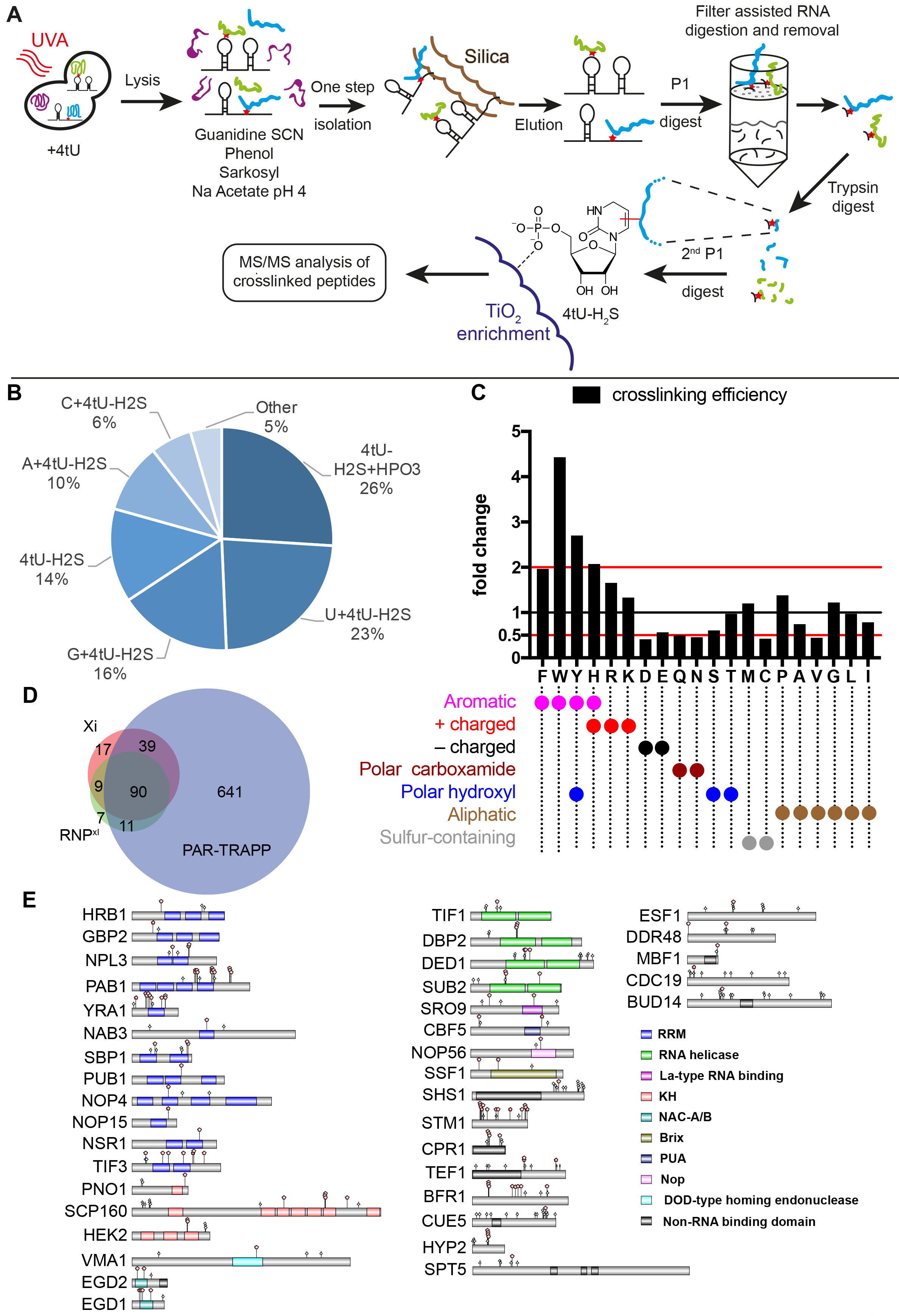
Identifying the RNA-crosslinked peptides with iTRAPP. (A) iTRAPP workflow to directly observe crosslinked RNA-peptides species by mass spectrometry. See the main text for details. (B) Pie chart of RNA species observed crosslinked to peptides by the Xi search engine run in targeted mode. (C) The analysis of amino acids, reported as crosslinked by the Xi search engine run in targeted search mode. Amino acids are represented by single letter IUPAC codes. Black bars-crosslink efficiency, defined as ratio between the frequency of crosslinked amino acids and the frequency of amino acids in all crosslinked peptides. (D) Venn diagram showing the overlap between proteins identified in PAR-TRAPP, RNP^xl^ and Xi. (E) Domain structure of selected proteins, identified as crosslinked by the Xi search engine. Domains (colored rectangles) and sites of phosphorylation (light green rhombi) from the uniprot database were plotted onto proteins represented by grey rectangles. Crosslink sites, identified by Xi are indicated with red pentagons. See also Supplementary Text.

**Figure 6.**
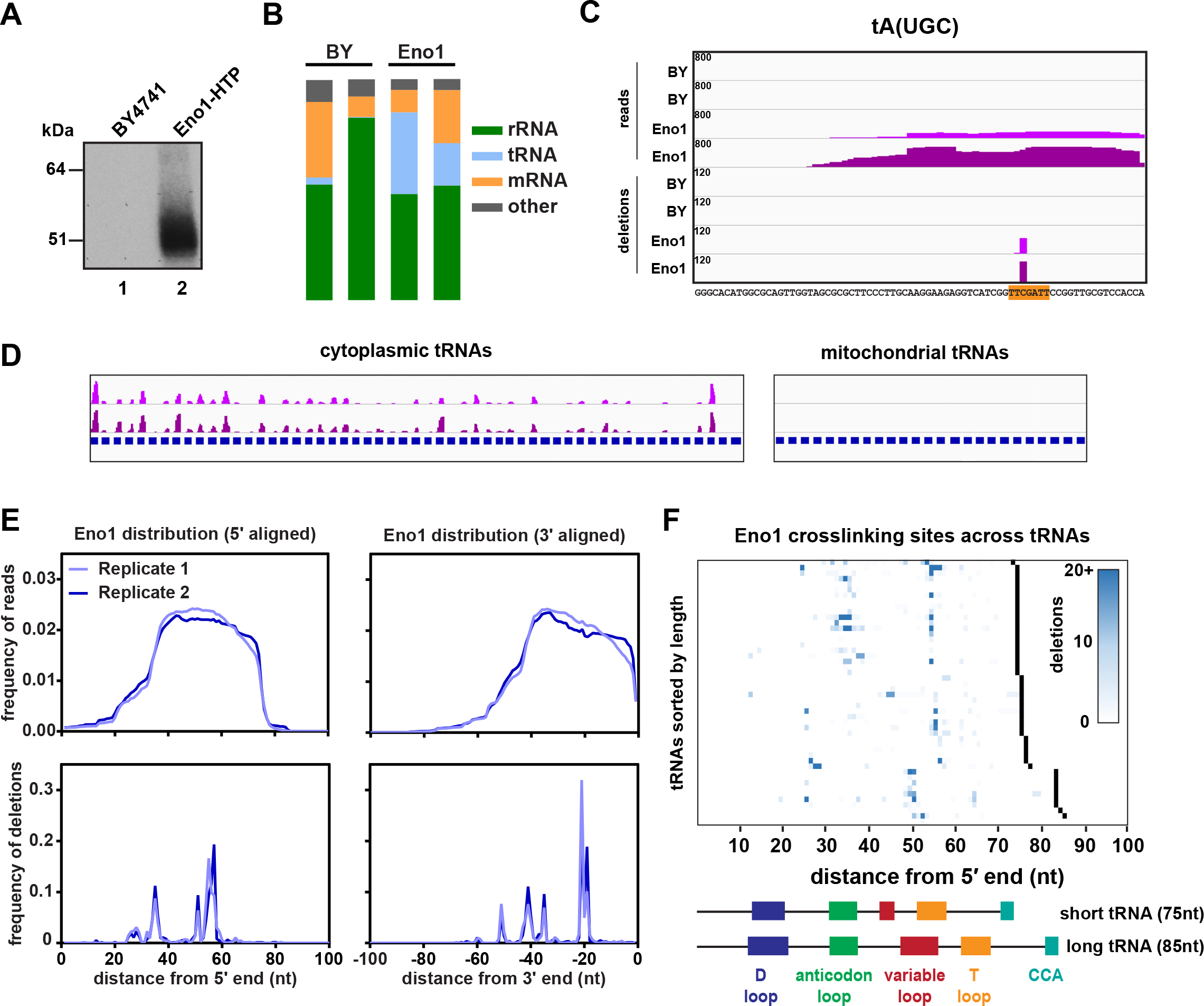
(A) Representative gel showing the recovery of radiolabeled RNA after CRAC purification. Lane 1: Untagged control strain (BY4741). Lane 2: Strain expressing Eno1-HTP from its endogenous locus. (B) Bar charts showing the relative distribution of reads among different classes of RNA. (C) The binding of Eno1 to the representative tRNA tA(UGC). The four upper tracks show the distribution of entire reads, while the four lower tracks show putative crosslinking sites (deletions). Tracks are scaled by reads (or deletions) per million, and this value is denoted in the upper left corner of each track. Two independent replicates are shown for the untagged BY control and Eno1-HTP. The tRNA sequence is shown below with the T-loop sequence highlighted in orange. (D) Global view of Eno1 binding to cytoplasmic tRNAs (left) or mitochondrial tRNAs (right). For ease of viewing, tRNAs across the genome were concatenated into a single ‘chromosome’, with each tRNA gene annotation shown in blue. Two independent replicates are shown. (E) Metagene plots showing the distribution of reads or deletions summed across all tRNAs. tRNA genes were aligned from either the 5′ end (left) or the 3′ end (right). (F) Heat map showing the distribution of putative crosslinking sites (deletions) across all Eno1-bound tRNAs. tRNA genes are sorted by increasing length, and the 3′ end for each gene is denoted in black. The domain structures of a typical short (tH(GUG)) and long (tS(AGA)) tRNA are included for comparison.

In previous analyses (Kramer et al., 2014), and our initial analyses using 254 nm UV irradiation (Peil, Waghmare, Fischer, Spitzer, Tollervey and Rappsilber, in preparation), the crosslinked nucleotide recovered was predominately uridine monophosphate. We therefore used 4tU labelling and UVA irradiation to generate RNA-protein complexes for iTRAPP analyses (Figure 5A). Purified material eluted from silica was treated with nuclease P1 to completely degrade bound RNA to 5′-monophosphates. To remove non-crosslinked nucleotides, proteins were denatured by addition of 8M urea and loaded onto a 30kDa cut-off filter, which removed of 99.4% of the nucleic acids initially present in the sample (~40mg). Denaturation at this step is essential to prevent proteins smaller than 30kDa from passing through the membrane (Wiśniewski et al., 2011). Proteins were recovered from the membrane and digested with trypsin, conjugated to a solid support. This allows removal of trypsin from the sample once protein digestion is completed, permitting a second nuclease P1 digestion to ensure robust RNA degradation without protein protection of the RNA. The obtained peptides were desalted and analyzed by mass spectrometry.

Interpreting the spectra of an RNA-conjugated peptide represents a major challenge due to the fact that both RNA and peptide fragment generate rich ion spectra. The most informative peptide fragments bear the fragmented RNA, but the mass shift requires their annotation by specialist search engines. We compared the results of interpreting spectra using the published RNP^XL^ pipeline (Veit et al., 2016) and the Xi search engine, which was designed for peptide-peptide crosslinks (Giese et al., 2015; Mendes et al., 2018) (https://github.com/Rappsilber-Laboratory/XiSearch). Better results were obtained with Xi (see Supplementary text), which was used for subsequent analyses. RNA-modifications were identified with XiSearch in a targeted-modification search mode (Table S7). Crosslinked nucleotides were treated as “cleavable molecules” during collision induced dissociation in the gas phase. We generated search lists of 4-thiouridine derived fragments allowed to persist on the peptide fragments upon fragmentation, without defining amino acid specificity (See Materials and Methods). We used the number of peptide to spectrum matches passing the 1% FDR cut-off to identify the optimal fragment list. With the best performing search parameters, we identified 750 unique RNA-peptide crosslinks with 395 different peptides belonging to 155 proteins (Table 1 and Table S8).

First, we analyzed the RNAs identified as crosslinked to proteins (Figure 5B). As previously observed (Kramer et al., 2011; Kramer et al., 2014) the bulk of crosslinked nucleotides contained 4-thiouridine, lacking the mass of SH_2_. Surprisingly, the most frequent RNA adduct was 4-thiouridine containing 2 phosphate groups (4tU-H_2_S+HPO_3_), while nuclease P1, used for RNA degradation, generates 5′ monophosphate nucleotides. This may be a result of degradation of dinucleotides during sample preparation, as the second and third most frequent RNA species found in iTRAPP were U+4tU and G+4tU dinucleotides.

We next calculated the crosslinking efficiency for each amino acid species. This was defined as the ratio between the abundance of the amino acid in the pool of crosslinked peptides and its frequency at the crosslink site. (Figure 5C). Xi identified tryptophan (W), tyrosine (Y), phenylalanine (F), arginine (R), histidine (H), proline (P) and lysine (K) as crosslinked to RNA more than their frequency in the crosslinked peptides would predict. The high quality spectra with unambiguous mapping of the crosslink site for amino acids with reduced crosslinking efficiency could still be obtained (Figures S7-S9), showing that the crosslinks are detectable, and reduced efficiency probably reflects probability of a given amino acid to RNA crosslink.

Thus, aromatic residues tend to have a clear preference for crosslinking, presumably due to stacking interactions with the bases. Positively charged amino acids, recognized by trypsin protease, also showed elevated crosslinking. The presence of a negatively charged RNA near trypsin cleavage site is expected to interfere with trypsin digest of the crosslinked protein. Indeed, 75.7% of crosslinked RNA-peptide crosslinks had 1 or 2 missed trypsin cleavages. This number rose to 97.6% for peptides in which Xi reported lysine or arginine as the crosslinked amino acid, indicating that trypsin cleavage is unlikely to occur at or near to the site of crosslinking.

Proteins identified by iTRAPP show significant, but not complete, overlap with PAR-TRAPP (Figure 5D). Out of 155 proteins with crosslinking sites mapped by Xi, 26 were not identified as significantly enriched in TRAPP. These included several characterized RNA-binding proteins (Table S9); e.g. rRNA methyltransferase Nop2, RNA surveillance factor Nab3 and mitochondrial ribosomal protein Pet123. As noted above, some proteins likely retain RNA binding activity even under denaturing conditions leading to poor UV enrichment, but these can be positively identified in iTRAPP.

Some 48% of the observed crosslinks overlapped with protein domains, previously implicated in RNA binding (figure 5E). We hypothesized that protein-RNA interactions might be regulated by post-translational modification. We therefore assessed the frequency with which RNA-protein crosslink sites are localized within 10 amino acids of a confirmed site of phosphorylation and ubiquitination, taken from Uniprot. RNA-protein crosslinks strongly correlated with sites of phosphorylation (P value 3.3×10^-6^) (Figure 5E), with weaker correlation to ubiquitination sites (P value 2×10^-2^).

### Eno1 binds RNA *in vivo*

Many metabolic enzymes showed greater recovery with TRAPP than PAR-TRAPP (see above). We therefore wanted to determine whether TRAPP-enriched factors are *bona fide* RNA binding proteins using a different technique. For this purpose, we selected the yeast glycolytic enzyme Eno1 (enolase), which was robustly identified by TRAPP, but showed enrichment of less than 2-fold in PAR-TRAPP. Enolase was also confidently identified as RNA binding by TRAPP in *E. coli*, as expected due to its presence in the RNA degradosome complex (Carpousis, 2007), and the homologue, Eno2, was previously reported to interact with mitochondrial tRNAs (Entelis et al., 2006).

To tag Eno1, the chromosomal *ENO1* gene was fused to a tripartite C-terminal tag, His6-TEV protease cleavage site-Protein A (HTP), and the fusion protein was expressed from endogenous *P_ENO1_*. The strain expressing only Eno1-HTP showed wild-type growth (data not shown) indicating that the fusion protein is functional. This strain was then used in CRAC analyses, alongside the non-tagged negative control strain. In CRAC, proteins are subject to multistep affinity purification that includes two fully denaturing steps.

Eno1-HTP was efficiently crosslinked to RNA in actively growing cells using 254 nm UVC. Figure 7A shows [5′ ^32^P] labelling of the RNA-associated with purified Eno1, relative to the background in a non-tagged strain. High-throughput sequencing of cDNAs generated from RNAs bound to Eno1 (Figure 7B and Table S10) showed a wide distribution of target classes.

Among the top mRNAs bound by Eno1, the only overall enrichment was for highly expressed genes (Supp. Table S10). The *ENO2* mRNA was ranked at position 3, suggesting the possibility of regulation. However, inspection of the hit distribution across the *ENO2* mRNA did not indicate any specific binding site, which might have been expected for a regulatory interaction. This was also the case for all other mRNAs identified with good sequence coverage. In contrast, the distribution of Eno1 binding appeared to show greater specificity on tRNAs, which comprised ~25% of the recovered sequences (Figure 7B). The structure and modifications present in tRNAs reduces their recovery by standard RT-PCR, so this recovery is likely to under-estimate the frequency of Eno1-tRNA interaction *in vivo*. Recovery was specific for cytoplasmic tRNAs relative to mitochondrial tRNAs (Figure 7D), showing that these reflect *bona fide*, *in vivo* interactions., with specific enrichment for binding to tRNAs.

Within tRNAs, Eno1 binding sites showed high specificity. For example, the crosslinking site in tRNA^Ala^ was localized to a single nucleotide within the T-loop (Figure 7C). Similar patterns were seen on many tRNAs, with Eno1 bound to either the T-loop, the anticodon loop, or both (Figure 7E-F). On a few, long tRNAs, Eno1 showed binding to the extended variable loop (Figure 7F). These results indicate that Eno1 binds to specific structural elements within cytoplasmic tRNAs and highlight the potential for TRAPP to uncover novel RNA interactions.

## DISCUSSION

The identification of ever-increasing numbers of RNA species has underlined the importance of robust characterization of *bona fide* sites of protein-RNA interaction. Here, we report the development and application of protocols for the purification of the RNA-bound proteome, providing quantitative data on protein crosslinking efficiencies under normal or stress conditions, and identifying precise sites of protein-RNA contact.

### Differences between TRAPP and PAR-TRAPP results

An unexpected finding was the differences in protein recovery in TRAPP using 254 nm UVC irradiation compared to PAR-TRAPP using 4-thiouracil incorporation combined with long-wavelength UVA irradiation. In both cases, the most abundant classes of proteins recovered correspond to expected targets, including the translation machinery and many other annotated RNA interacting proteins. However, there were clear differences in the recovery of proteins that lack known functions in RNA metabolism, which were much more abundant in TRAPP. In particular, metabolic enzymes were major targets in TRAPP, including almost all glycolytic enzymes. There is every reason to think that these data capture genuine, *in vivo* RNA-protein contacts. In our data, the protein recovery is clearly significant and reproducible, and for Eno1 we confirmed robust and specific *in vivo* crosslinking. Moreover, similar interactions have been independently identified in other analyses of RNA-binding proteomes using UVC irradiation (Baltz et al., 2012; Castello et al., 2012; Kramer et al., 2014; Kwon et al., 2013). Strikingly, there was strong conservation of RNA-binding between homologous enzymes of intermediary metabolism yeast and *E. coli*, which is generally a sign of a significant, conserved function.

The basis of the difference between crosslinking with 254 nm UV compared to 4-thiouracil with UVA may lie in the properties of the nucleotides. Native nucleotides show very rapid UV radiation decay with subpicosecond lifetimes (i.e. <10^-12^ s) (reviewed in (Beckstead et al., 2016)). This likely reflects non-radiative decay of the activated state, with the energy dissipating as heat, protecting the nucleotide from damage. This may be a feature that was selected early in the development of life, presumably in a very high UV environment (Beckstead et al., 2016). We speculate that during 254 nm irradiation, nucleotides can absorb many photons but will normally very rapidly release the energy as heat. However, if in close contact with an amino acid, crosslinking can occur during this short time window. This potentially allows multiple opportunities for crosslinking with transiently interacting proteins, although the frequency will be low for any individual nucleotide. In contrast, 4-thiouridine absorbs UVA and generates a reactive triplet with high quantum yield (0.9) (Zou et al., 2014). The 4-thiouridine can be crosslinked to an amino acid, if suitably positioned at that instant. Alternatively, a highly reactive singlet oxygen can be generated, which reacts with 4-thiouridine to generate uridine or uridine sulfonate (U-SO_3_). In either case, no further crosslinking can occur. Our interpretation of the difference between the TRAPP and PAR-TRAPP data is, therefore, that 254 nm UV allows multiple chances of crosslinking during the period of irradiation, increasing the probability of capturing weak transient interactions. In contrast, 4-thiouridine reacts with UV only once and is therefore biased towards the recovery of stable interactions. This hypothesis leads us to propose that the many proteins specifically identified by TRAPP that lack known RNA-related functions and were not significantly enriched by PAR-TRAPP, will generally form weak, transient RNA interactions.

### A potential role for RNA-protein interactions in modulating phase-separation

There has been substantial recent interest in the phenomenon of liquid-liquid phase separation or demixing (reviewed in (Boeynaems et al., 2018)). This is based on observations that at high concentrations, many specific proteins or protein-RNA complexes can spontaneously self-assemble into liquid-like droplets under suitable conditions *in vitro* (Lin et al., 2015). This phenomenon is proposed to underpin the formation of multiple, non-membrane bound, subcellular structures – particularly those involving RNA-protein complexes, including the nucleolus, nuclear Cajal bodies or cytoplasmic P-bodies and stress granules. This model rationalizes long standing observations that these structures show liquid-like properties (see, for example (Wu et al., 1991)) and spontaneously assemble due to multiple weak interactions between the components (reviewed in (Lewis and Tollervey, 2000)). Different RNAs, proteins and RNP complexes can be greatly concentrated by these processes, presumably enhancing the efficiency of RNP assembly and processing. RNA interactions help drive phase separation in many cellular contexts, e.g. stress granule formation nucleated by free mRNAs and paraspeckles nucleated by NEAT1 ncRNA (Clemson et al., 2009; Lin et al., 2015). This may be opposed or regulated by low affinity, heterogeneous protein binding. In addition, RNA interactions regulate the behavior of prion-like proteins, with reduced RNA binding associated with increased phase separation and the formation of cytotoxic solid-like assemblies (Maharana et al., 2018). We speculate that the anti-phase separation property of RNA is utilized by the abundant metabolic proteins for which uncontrolled self-assembly into separated cellular domains seems highly undesirable. The concentration of these proteins in the cell is very high, so inappropriate phase separation may pose a substantial potential problem. Since phase separation is driven by self-assembly, it might be anticipated that it would be opposed by the formation of large numbers of different, non-self-interactions with abundant structured RNA species. We speculate that this underpins the tendency of well-structured proteins to have domains or sites that form multiple RNA interactions. Furthermore, lack of such correlation in PAR-TRAPP, demonstrated in Figure 1G, suggests that most of the unconventional RNA binding proteins analyzed in Figure 1G interact very transiently with cellular RNAs to avoid getting trapped into one of the variety of phase separated granules.

We postulate that many of the novel RNA-protein interactions do not have specific, pair-wise functional roles, but rather act biophysically “*en masse*” to modulate cell organization. This does not, of course, preclude the possibility that individual, novel RNA-protein interactions do play specific and important roles. If this an important property, the ability of enzymes of intermediary metabolism to interact with RNA should be conserved through evolution. Indeed, this was observed for comparisons of *S. cerevisiae* and *E. coli* RNA-interacting proteome, where the same conserved enzymes, catalyzing the same reactions were observed interacting with RNA. Thus, we believe that further investigation into the role of RNA-interacting metabolic enzymes in stress response is warranted.

To test this model, we analyzed the major glycolytic enzyme Eno1, which gave substantially stronger crosslinking in TRAPP than in PAR-TRAPP. This revealed preferential interaction of Eno1 with the 3′ regions of cytoplasmic tRNAs, which are suitable RNAs to interact with to achieve an anti-phase separation effect. Our data suggests that each tRNA offers one major binding site for Eno1 protein. In addition, tRNA molecules are highly structured and are thus unlikely to participate in RNA-RNA interactions that might promote phase separation. Notably, several other glycolytic enzymes were reported to form phase separated, cytoplasmic “G bodies” following hypoxic stress (Jin et al., 2017; Miura et al., 2013). It seems possible that formation of these bodies might be regulated by protein-RNA interactions.

### PAR-TRAPP identifies stress-induced changes in RNA-protein interactions

We took advantage of the ease and reproducibility of TRAPP approaches to analyze the immediate effects of weak acid stress on the RNA-bound proteome. Analyses were performed comparing non-stressed cells, to cells treated with sorbic acid for 16 min, to reduce the intercellular pH. Yeast generally grows in acid conditions and reduced intracellular pH is a common feature in response to multiple stresses, including glucose deprivation, due to inactivation of the membrane proton pumps that maintain the pH gradient. Moreover, reduced pH was reported to solidify the cytoplasm, due to charge neutralization on abundant proteins (Munder et al., 2016). We speculated that this might also be associated with altered RNA interactions by cytoplasmic proteins.

Only around 10% of RNA-interacting proteins showed differences in crosslinking of greater than 2-fold, suggesting that the stress responses are relatively specific. This was supported by analyses of individual pathways of RNA metabolism. For example, ribosome biogenesis, which is expected to be inhibited by stress, appeared to be blocked at a specific step in the cytoplasmic maturation of the precursors to 60S subunits.

Multiple proteins involved in translation showed reduced interaction following sorbic acid treatment, presumably reflecting general translation inhibition. Less expectedly, these included factors reported to be P-body components; eiF4G (Tif4631 and Tif4632), the RNA helicase Ded1, and the poly(A) RNA binding proteins Pub1 and Pab1. Moreover, P-body components directly implicated in mRNA degradation also showed reduced RNA interactions; including the CCR4/NOT complex subunit Not3, decapping cofactors Pat1 and Edc3, surveillance factors, Upf3, Nmd2 and Ebs1, and the 5′ exonuclease Xrn1. In contrast, increased RNA interactions were observed for other P-body-associated proteins: These included the translation repressor Ssd1 and the dissociable RNAPII component Rpb7, as well as seven enzymes of intermediary metabolism, previously identified as P-body components. Increased RNA interaction upon sorbic acid exposure was also found for a further for additional 29 proteins of intermediary metabolism.

We speculate that stress signaling allows abundant proteins of intermediary metabolism to form more stable interactions with RNA, allowing PAR-TRAPP detection. RNA-protein interactions can facilitate liquid-liquid phase separation into droplets (Lin et al., 2015), which build P-bodies, stress granules and other phase separated regions, while RNA-RNA interactions may help guide the RNA species destined for the forming phase-separated granules (Langdon et al., 2018). In the short term this may provide protection, however, prolonged stress would potentially allow droplets to mature into more filamentous states with negative consequences (Lin et al., 2015).

### Mapping precise amino acid-RNA contacts with iTRAPP

A previous report (Kramer et al., 2014) and our initial observations indicated that adenosine, guanosine and cytosine are very poorly represented in RNA-protein crosslinks that can be identified by MS. The basis for this is unclear, since reciprocal CRAC and iCLIP mapping of interaction sites on the RNA shows no such extreme bias. However, we took advantage of this finding and developed the iTRAPP protocol based on 4tU labelling. This provides greater specificity for RNA-binding proteins than 254 nm crosslinking, with reduced complexity of RNA crosslinks since only 4-thioridine containing species were expected.

Most amino acids can form photo-adducts with oligo-DNA or RNA sequences when single amino acids are added to the reaction (see for example (Shetlar, 1980). In our analyses, crosslinks were identified with all amino acids, with over-representation of aromatic amino acids (Trp, Tyr, Phe) relative to their abundance in the crosslinked peptides, followed by positively charged amino acids (His, Lys, Arg). Less expectedly, the hydrophobic amino acids (Met, Gly, Pro) were also enriched, possibly reflecting their role in unstructured domains (van der Lee et al., 2014).

Despite progress in the software to analyze RNA-peptide crosslinks much remains to be done in this field. The chemistry of RNA-protein crosslinks is more complicated than just a sum of RNA and peptide masses. For example, hydrogen losses were demonstrated in the case of lysine crosslinking to 4-thiouridine (Kramer et al., 2011). The challenge is that allowing the flexibility required to identify all possible crosslinks and fragmentation products, greatly increases the number of false positive hits to a decoy database. Better understanding of RNA-protein crosslink chemistry would allow the development of software that is aware of specific predicted mass losses or gains depending on which amino acid participates in the crosslinking.

Comparison of precise sites of protein-RNA contact to published sites of protein modification revealed highly significant enrichment for proximity to reported sites of protein phosphorylation and ubiquitination. We predict that these modifications will negatively regulate closely positioned RNA-protein interactions, by charge and/or steric hindrance. This could be defined by future analyses combining TRAPP with mutations in protein modifying enzymes.

The protocol described here can be readily adapted to other species. We anticipate that the resources provided here, a protocol to study protein-RNA binding sites at global scale and a list of protein-RNA attachment sites in *S. cerevisiae*, will stimulate systematic and detailed studies of the functionally important protein-RNA interface.

## MATERIALS AND METHODS

### Yeast SILAC media and crosslinking

Yeast cells (BY4741 derived MATa, *his3*Δ*1 leu2*Δ*0 met15*Δ*0 ura3*Δ*0, arg4*Δ, *Lys9*Δ) were grown at 30°C under shaking in yeast nitrogen base media (Formedium CYN0410), containing 2% w/v of D-Glucose (Fisher G/0500/60), supplemented with Complete Supplement Mixture without tryptophan, lysine, arginine and uracil (Formedium DCS1339) and with 20 mg/L Uracil (Sigma U0750-100g). SILAC light media was additionally supplemented with 30 mg/L lysine (Sigma L5626-100g) and 5 mg/l arginine (A5131-100g), while 30mg l^-1^ 13C6 lysine (Cambridge Isotope Laboratories CLM-2247-H-0.25) and 5mg l^-1^ 13C6 arginine (Cambridge Isotope Laboratories CLM-2265-0.5) was added to SILAC heavy media. Equal cell mass (by OD measurement) of Heavy-labelled and light-labelled cells were combined per experiment.

For TRAPP using UVC (254nm) crosslinking, the culture was harvested at OD_600_ 0.5 from 750ml of media and irradiated with Megatron apparatus (UVO3) as described (Granneman et al., 2011). Cells were harvested by filtration and immediately frozen for later processing.

For PAR-TRAPP, using UVA (350 nm) crosslinking, cells were grown to OD 0.15, 4-thiouracil (4tU) (Sigma 440736-1g), was added to 0.5 mM from 1 M DMSO stock solution and cells were allowed to grow for 3 hours. Cells were harvested by filtration and resuspended in 800ml of growth media without 4tU cells were placed on the irradiation tray of the eBox irradiation apparatus. The UVA lamps were pre-warmed for 1min, and the cells were irradiated for 38sec delivering 7.33J cm^-2^ of UVA. Cells were harvested by filtration and frozen for later processing.

### *E. coli* SILAC media and crosslinking

*E. coli* cells (MG1655 derived F-, lambda-, rph-1 ΔlysA ΔargA) were grown in 150 ml of M9 minimal medium, supplemented with 1μM FeNH_4_SO_4_, 0.2% w/v of D-Glucose (Fisher G/0500/60), 1 mM MgSO_4_. SILAC light media was additionally supplemented with 40mg/L lysine (Sigma L5626-100g) and 40mg l^-1^ arginine (A5131-100g), while 40mg l^-1^ [^13^C_6_]-lysine (Cambridge Isotope Laboratories CLM-2247-H-0.25) and 40mg l^-1^ [^13^C_6_]-arginine (Cambridge Isotope Laboratories CLM-2265-0.5) was added to SILAC heavy media. Cells were grown to OD 0.5, diluted to 700 mL with growth media and irradiated with UVC (254 nm) using a Minitron (UVO3) (Granneman et al., 2011). Equal cell mass (70 ODs) of Heavy-labelled and light-labelled cells were combined per experiment.

### Sorbic acid treatment

Yeast cells were treated as described above with 4tU in SILAC yeast nitrogen base media for 3 hours. Sorbic acid was added to 6mM final concentration from ethanol stock solution and cells were incubated for 16min. Cells were harvested and irradiated as described above.

### eBOX irradiation apparatus

The e-box is a chamber with internal dimensions of 49.5×25.5×48 cm (W×H×D) lined with aluminium foil. The sample tray made of borosilicate glass is positioned between 2 banks of 10 PL-L 36W/09/4P PUVA 350 nm bulbs (Philips 871150061410040). Each bank is driven by 5 PC236TCLPRO-TR ballasts (Tridonic 22176170). The two banks together consume 720 W of electrical power. UVA measurements were performed with Sper scientific UVA/B Light Meter (850009), equipped with UVFS Reflective ND Filter, OD: 2.0 (THORLABS NDUV20B). Based on the spectral sensitivity of the meter, we estimate that in 38 seconds of irradiation time we administer from 7.3 J cm^-2^ of UVA (193mW cm^-2^). The UVA output of the eBox stabilizes after 1min of lamp warmup. The sample is prevented from UVA exposure with sliding shutters during the warmup period, followed by shutters extraction and sample irradiation for 38sec.

### Silica bead preparation

Silica beads (Fluka S5631-500g) were resuspended in 1M HCl and left for 24h at room temperature. The beads were then washed with water 3 times by spinning in a 50ml falcon tube at 3000 rcf for 2min and decanting the supernatant, containing beads of the smallest size. After the last wash beads are resuspended in volume of water to achieve 50% slurry suspension.

### Cell lysis

*E. coli* cells pellets (140 ODs) were resuspended in 8ml of 1:1 mix between Phenol (Sigma P4557), saturated with 10mM Tris-HCl pH 8.0, 1mM EDTA and GTC buffer (4 M Guanidine Thiocyanate (Fisher BP221-1), 50 mM Tris-HCl pH 8.0 (Invitrogen 15504-020), 10 mM EDTA (Fisher D/0700/53), 1% betamercaptoethanol), sample was split between 8 2ml screw-cap tubes, 1 volume of zirconia beads (BioSpec 11079105z) were added and cells were lysed with FastPrep-24 5g (MP biomedicals 116005500) twice for 40sec at 6 m sec^-1^. Lysates were recovered into 50ml falcon tube.

Yeast mixed cells pellets were resuspended in 2ml of Phenol-GTC buffer in a 50ml falcon tube 3ml of Zirkonia beads were added and cells were vortexed for 1min 6 times with 1min incubation on ice between vortexing. 6ml of phenol-GTC mix was added and lysates were vortexed for additional 1 min.

### RNP isolation

Samples in 50ml falcon tubes were spun down at 4600 rcf, 4°C for 5min. Supernatant was recovered into 2ml tubes and the samples were spun at 13000 rcf for 10min. The supernatant was recovered into a 50ml tube, 0.1 volume of 3M Sodium Acetate-Acetic acid pH 4.0 was added, followed by 1 volume of absolute ethanol. The sample was mixed by brief vortexing and 1ml of 50% slurry suspension of Silica was added. Nucleic acids were allowed to load onto silica for 60 minutes on a rotating wheel at room temperature. The beads were spun down at 2500 rcf, 4°C for 2min. The beads were resuspended by vigorous vortexing in 15 ml of wash buffer I (4M Guanidine Thiocyanate, 1 M Sodium acetate (Alfa Aesar 11554) - Acetic acid (Fisher A/0400/PB17) pH 4.0, 30% Ethanol) and spun again at 2500 rcf, 4°C for 2min. The supernatant was discarded and the wash was repeated 2 more times. Finally, the beads were washed 3 times with 20ml of wash buffer II (100mM NaCl, 50mM Tris-HCl pH6.4, 80% Ethanol). After the last wash, the beads were resuspended in a small amount of wash buffer II, and transferred to 2 2ml tubes. The tubes were spun at 2000 rcf for 2min at RT and the supernatant was discarded. Silica sand was dried for 10min in a speedvac at 45°C. Nucleic acids were eluted 3 times with 500μl. 1μl of RNaseA+T1 (Ambion AM2286) mix was added and the samples were incubated for 2 hours in a speedvac at 45°C, followed by speedvac incubation at 65°C until the samples are fully dry.

The dried samples are resuspended in 75 μl Laemli buffer by vortexing, incubated at 100°C for 10min. 10μl were resolved on a gradient 4-20% Mini-PROTEAN TGX gel (Biorad 4561095), stained with Imperial protein stain (Thermo 24615) according to the manufacturers protocol. This gel is either processed further or is used to estimate the amount of material per sample.

Each lane of the gel, corresponding to one SILAC mix was cut into at least 4 gel fractions. Excised fractions were de-stained and proteins were digested with trypsin, as previously described (Shevchenko et al., 1997). Briefly, proteins were reduced in 10mM dithiothreitol (Sigma Aldrich, UK) for 30min at 37°C and alkylated in 55mM iodoacetamide (Sigma Aldrich, UK) for 20 min at ambient temperature in the dark. They were then digested overnight at 37°C with 12.5ng μl^-1^ trypsin (Pierce, UK).

Following digestion, samples were diluted with equal volume of 0.1% TFA and spun onto StageTips as described (Rappsilber et al., 2003). Peptides were eluted in 20μl of 80% acetonitrile (ACN) in 0.1% trifluoroacetic acid (TFA) and concentrated down to 4μl by vacuum centrifugation (Concentrator 5301, Eppendorf, UK). The peptide sample was then prepared for LC-MS/MS analysis by diluting it to 5μl by 0.1% TFA.

Where indicated, treatment of nucleic acids with DNase, was performed in buffer containing Tris pH 8.0, 2.5 mM MgCl_2_, 0.5 mM CaCl_2_, using DNase I from New England Biolabs (M0303) at 100U/mg of DNA, supplemeted with 200U of RNAsin (Promega N211A). Treatment with Cyanase (Serva 18542.01) was performed in Tris pH 8.0 buffer supplemented with 6mM MnSO_4_, using 0.1U of enzyme per 1μg of nucleic acids. Silver staining was performed as previously described in (Yan et al., 2000).

### iTRAPP

Yeast cells were with UVA as described above. Altogether 10L of cells were processed as one sample. Cells were lysed and RNA associated protein was isolated as described above. After RNA elution from silica beads, sodium acetate pH5.3 was added to 10mM ZnCl_2_ to 0.5mM and DTT to 0.5 mM final concentrations. Nuclease P1 (sigma N8630) was added 1:4000 weight of RNA: weight of enzyme and the sample was incubated at 37°C overnight. Tris-HCl pH8.0 buffer was added to 50 mM, DTT to 30mM and solid urea (Acros organics 327380050) was added for 8 M final concentration to reduce and denature proteins. The solution was passed through a 30KDa MWCO Vivaspin Polyethersulfone concentrator (GE Healthcare 28-9323-61) and flow through was discarded. Retentate was washed 3 times with urea buffer (8M urea in 50mM Tris-HCl pH8.0) and cysteines were blocked by passing through a solution of 50 mM IAA in urea buffer. The retentate was washed 3 times with 2mM DTT in urea buffer to quench the IAA. Finally, retentate was recovered, urea was diluted to 1M with 50mM Tris pH8.0. The sample was added to trypsin, conjugated with magnetic beads (Takara 635646) and the sample was incubated for 12h with mixing at 37°C. Next, trypsin beads were removed from the supernatant and the sample was diluted with water 1:1. Sodium acetate was added to 10mM, ZnCl_2_ to 0.1mM and pH was reduced to approximately 5.2 with acetic acid. 20μg of nuclease P1 was added and the sample was digested at 37°C for 24h.

TFA was added to 1%, ACN to 2.5% final concentration. Sample was loaded onto 4 C18 SPE cartridges (Empore 4215SD). Washed with 0.1% TFA and eluted in 0.1% TFA, 80% ACN.

Eluates were dried in speedvac. Next the sample was resuspended in 250μl of binding buffer (50% ACN, 2 M lactic acid) with vortexing and brief sonication. Phospho group containing RNA-crosslinked peptides were the captured with TiO_2_ Mag Sepharose beads (GE healthcare 28- 9440-10) according to the manufacturers protocol. Peptides eluted from TiO_2_ sepharose were loaded onto C18-StageTtips as described above followed by LC-MS analysis.

### LC-MS analysis

LC-MS analysis were performed on an Orbitrap Fusion™ Lumos™ Tribrid™ Mass Spectrometer (Thermo Fisher Scientific, UK) coupled on-line, to an Ultimate 3000 RSLCnano Systems (Dionex, Thermo Fisher Scientific, UK). Peptides were separated on a 50 cm EASY-Spray column (Thermo Scientific, UK), which was assembled on an EASY-Spray source (Thermo Scientific, UK) and operated at 50°C. Mobile phase A consisted of 0.1% formic acid in LC-MS grade water and mobile phase B consisted of 80% acetonitrile and 0.1% formic acid. Peptides were loaded onto the column at a flow rate of 0.3μl min^-1^ and eluted at a flow rate of 0.2 μl min^-1^ according to the following gradient: 2 to 40% mobile phase B in 136min and then to 95% in 11min. Mobile phase B was retained at 95% for 5 min and returned back to 2% a minute after until the end of the run (160 min). The iTRAPP analysis was performed on a QExactive mass spectrometer (Thermo Fisher Scientific, UK). The same LC conditions applied as described above.

For orbitrap Fusion Lumos, FTMS spectra were recorded at 120,000 resolution (scan range 400-1900 m/z) with an ion target of 4.0e5. MS2 in the ion trap at normal scan rates with ion target of 1.0E4 and HCD fragmentation (Olsen et al., 2007) with normalized collision energy of 27. The isolation window in the quadrupole was 1.4 Thomson. Only ions with charge between 2 and 7 were selected for MS2.

For QExactive, FTMS spectra were recorded at 70,000 resolution and the top 10 most abundant peaks with charge ≥ 2 and isolation window of 2.0 Thomson were selected and fragmented by higher-energy collisional dissociation with normalized collision energy of 27. The maximum ion injection time for the MS and MS2 scans was set to 20 and 60ms respectively and the AGC target was set to 1 E6 for the MS scan and to 5 E4 for the MS2 scan. Dynamic exclusion was set to 60s.

The MaxQuant software platform (Cox and Mann, 2008) version 1.6.1.0 was used to process the raw files and search was conducted against the *S. cerevisiae* reference proteome set of Uniprot database (released on November 2017), using the Andromeda search engine (Cox et al., 2011). The first search peptide tolerance was set to 20ppm while the main search peptide tolerance was set to 4.5ppm. Isotope mass tolerance was 2ppm and maximum charge to 7. Maximum of two missed cleavages were allowed. Carbamidomethylation of cysteine was set as fixed modification. Oxidation of methionine, acetylation of the N-terminal and phosphorylation of serine, threonine and tyrosine were set as variable modifications. Multiplicity was set to 2 and for heavy labels Arginine 6 and Lysine 6 were selected. Peptide and protein identifications were filtered to 1% FDR. For the iTRAPP experiment, peak lists were generated from Max Quant version 1.6.1.0. The Xi software platform was used to identify proteins cross-linked to RNA.

### TRAPP UV enrichment data analysis

Peptides.txt file, generated by maxquant software was used for the analysis. First, peptides mapping to contaminants and decoy database were removed. The reported heavy and light intensities per peptide were exported into Rstudio version 1.1.453 (http://www.rstudio.com/). Missing peptide intensity values were imputed using impute.minprob function of the imputeLCMD package version 2.0 (Lazar et al., 2016) (https://rdrr.io/cran/imputeLCMD/). For *S. cerevisiae* PAR-TRAPP and *E. coli* TRAPP data we run impute.minprob function with the following parameters: q=0.1 tune.sigma=0.01. For *S. cerevisiae* TRAPP data tune.sigma parameter was set to 0.0035. If in a given biological replicate a peptide was missing intensity in both + UV and −UV, the imputed values were removed. We then calculated +UV to −UV protein ratio in each experiment as median of peptides using "leading razor protein" as protein identifier. Proteins identified with at least 2 peptides in at least 2 experiments were considered for further analysis. The protein ratios were log_2_ transformed and the statistical significance of the result was determined using the Limma package version 3.36.2 (Ritchie et al., 2015).

### iTRAPP data analysis

The targeted Xi search was performed with cleavable RNA modifications limited to up to tri-nucleotide RNA, where at least one residue is 4tU (Table S7). The targeted masses can be identified on top of the otherwise defined variable modifications. Note that only one of the specified masses can be identified per PSM to prevent very long search times. Since the crosslink was assumed to always happen through the 4tU residue, the masses allowed to remain on fragment peptide ions were derived from 4tU: 128.004435 (4tU base); 94.016713 (4tU base-H2S); 226.058971 (4-thiouridine monophosphate-HPO3-H2S); 306.025302 (4-thiouridine monophosphate-H2S); 340.013027 (4-thiouridine monophosphate); 260.046696 (4-thiouridine monophosphate-HPO3); 385.991636 (4-thiouridine monophosphate+HPO3-H2S); 419.979358 (Uridine monophosphate+HPO3).

FDR was calculated on a list of peptide spectral matches, sorted by descending score, as a ratio between the number of decoy hits and the number of target hits for a given score. Xi search setup: The raw data from QExactive mass spectrometer was processed into peak lists using Maxquant 1.6.1.0. We then performed a search using Xi version 1.6.731 (https://github.com/Rappsilber-Laboratory/XiSearch) (Mendes et al., 2018) search against the reference proteome with the following search parameters:

MS1 tolerance 6 ppm; MS2 tolerance 20ppm; enzyme – trypsin\p; missed cleavages allowed – 2; minimum peptide length – 6; Fixed modifications - Carboxymethyl at cysteine; Variable modifications – oxidized methionine; maximum number of neutral losses per MS2 fragment – 1; Report MS2 fragments that are off by 1 Da – True; Complete search configuration can be found in the supplementary file 1.

RNP^xl^ plugin: We utilized RNP^xl^ plugin (Veit et al., 2016) for Proteome discoverer version 2.1.1.21 (Thermo Fisher Scientific). MS1 tolerance 6ppm; MS2 tolerance 20ppm; enzyme – trypsin\p; missed cleavages allowed – 2; Maximum RNA length – 3 nucleotides of which one has to be 4tU. RNA modifications - HPO3-H2S; -H2S; +HPO3-H2S; +HPO3;-HPO3; Fixed modifications - Carboxymethyl at cysteine; Variable modifications – oxidized methionine; In order to be able to calculate FDR for RNP^xl^ hits, we constructed a combined target + decoy database using Decoy database builder software (Reidegeld et al., 2008). The software was run in combined mode (reverse + shuffled + random decoy database). FDR was calculated on a list of peptide spectral matches, sorted by descending score, as a ratio between the number of decoy hits and the number of target hits for a given score.

### Scoring for co-localization of RNA-protein crosslinks with sites of post translational protein modifications

The modification sites for proteins recovered in iTRAPP were obtained from Uniprot database on 12.07.2018 (The UniProt Consortium, 2017). Peptide to spectrum matches passing the 1% FDR were processed in MS Excel to remove duplicates based on protein and crosslinked amino-acid (as reported by Xi). The list was further filtered to remove duplicates with same peptide and same RNA. Each filtering step was performed so as to keep peptide to spectrum matches with highest match score. We then calculated the number of observed crosslinks within 20 amino acids from a phospho site (X-Pi^observed^) and the number of observed crosslinks outside the 20 residue window (X^observed^). In order to obtain the expected values, we shuffled the phospho sites using the MS excel “RANDBETWEEN” function in the crosslinked proteins and calculated the (X-Pi^expected^) and (X^expected^). We performed the shuffling 100 times and calculated the average (X-Pi^expected^) and (X^expected^) values, Finally, we performed a chi-square test to identify if the observed values were significantly different from the averaged expected values. The analysis for ubiquitination sites was performed similarly, however we analyzed ubiquitination and sumoylation sites together as one modification.

### Bioinformatics resources

We utilized The Yeast Metabolome Database (Ramirez-Gaona et al., 2017) and The *E. coli* Metabolome Database (Sajed et al., 2016) to identify proteins involved in intermediary metabolism. The analysis of enrichment of proteins belonging to the GO terms in figures 1B, 1C, 2B, 2C, 3A and 4A was performed with Perseus software (Tyanova et al., 2016). GO term annotations for the analysis were downloaded from annotations.perseus-framework.org on the 29.11.2017 for *S. cerevisiae* and on 25.02.2018 for *E. coli*. The analysis shown in figures 1E, 1F, 3B, 3D, S5-S6 was performed using David 6.8 web service (Jiao et al., 2012). Venn diagrams were generated using the BioVenn web service (Hulsen et al., 2008). To identify orthologous clusters between *E. coli* and *S. cerevisiae* we took advantage of the Inparanoid 8 database (Sonnhammer and Östlund, 2015). Protein abundance data was obtained from the PaxDb database (Wang et al., 2015). Integrated dataset for S. cerevisiae was used for the analysis. The ratio between enriched and total proteins in Figure 1G was calculated by dividing the number of proteins in the bin annotated as UV enriched divided by the number of enriched proteins + the number of non-enriched proteins. The non-enriched proteins are defined as proteins quantified with 2 or more peptides in at least two experiments, but which fail to score a statistically significant enrichment upon UV irradiation (adjusted p value higher than 0.05).

### Sequencing data analysis

#### Pre-processing

Raw datasets were demultiplexed according to barcode using a custom python script. Flexbar (Dodt et al., 2012) was used to remove adaptor sequence, trim low quality bases from the 3′ end, and remove reads with low quality scores (parameters −u 3 -at 1 -ao 4 -q 30 with adapter sequence TGGAATTCTCGGGTGCCAAGGC). In addition to the barcode, each read contained three random nucleotides to allow PCR duplicates to be removed by collapsing identical sequences with fastx_collapser.

#### Alignment

Mapping sequencing reads to tRNAs is complicated by the fact that most tRNAs are present in multiple identical copies throughout the genome. Furthermore, mature tRNAs differ substantially from the genomic sequence: all tRNAs receive a CCA trinucleotide at the 3′ end, and some transcripts are posttranscriptionally spliced. For these reasons, sequencing reads were aligned to a database consisting of only mature tRNA sequences. A fasta file containing mature tRNA sequences was downloaded from Ensembl, and identical tRNAs were collapsed into single genes using fastx_collapser. Subsequently, the tRNA genes were concatenated into a single ‘chromosome’ with 40 base pair spacers between each tRNA using a custom python script. Similar files were also made for mRNAs and ribosomal RNAs. CRAC reads were mapped against this artificial genome using Novoalign (V2.07.00, Novocraft) and parameters −s 1 −l 17 – r None (to ensure only uniquely mapping reads). The accompanying Novoindex file was generated using parameters −k 10 −s 1. To remove PCR duplicates that were not previously discarded during pre-processing because of sequencing errors or differential trimming at the 3′ end, any reads with the same random nucleotides and with 5′ ends mapping to the same coordinates were collapsed into a single read (Tuck and Tollervey, 2013).

#### Downstream analysis

Downstream analyses were performed using the pyCRAC software (Webb et al., 2014). To count overlap with features, pyReadCounters (pyCRAC) was used together with a custom annotation file corresponding to the concatenated genome. Plots showing binding across tRNAs (Figures 7C-D) were generated with pyGTF2bedGraph (pyCRAC) using the results from pyReadCounters as input. The coverage at each position along the genome was normalized to the library size using the −permillion option. Putative crosslinking sites (deletions) were mapped using the −t deletions option. The heatmaps in Figure 7E were generated using the -outputall option in pyBinCollector and then plotted in Excel.

#### Data availability

All sequence data are available from GEO: accession number GSE119867.

The mass spectrometry proteomics data have been deposited to the ProteomeXchange Consortium via the PRIDE (Vizcaíno et al., 2016) partner repository with the dataset identifier PXD011071. Reviewer account details: Username; Password: lW5pwk7E.

## Author Contributions

V.S. and D.T. designed research; V.S. and E.P. performed biochemistry; C.S. and V.S. and performed and analyzed MS data, C.S., V.S. and L.F. performed bioinformatics; and V.S., S.B., C.S., J.R. and D.T. analyzed data and wrote the paper.

## Acknowledgements

VS was funded by the Swiss National Science Foundation (P2EZP3_159110). The Wellcome Trust generously funded this work through a Principle Research Fellowship to D.T. (077248), a Senior Research Fellowship to J.R. (103139) and an instrument grant (108504), with additional support from the Wellcome Trust/Edinburgh University via the Institutional Strategic Support Fund. Work in the Wellcome Centre for Cell Biology is supported by a Centre Core grant (203149).

---

**Figure S1.**
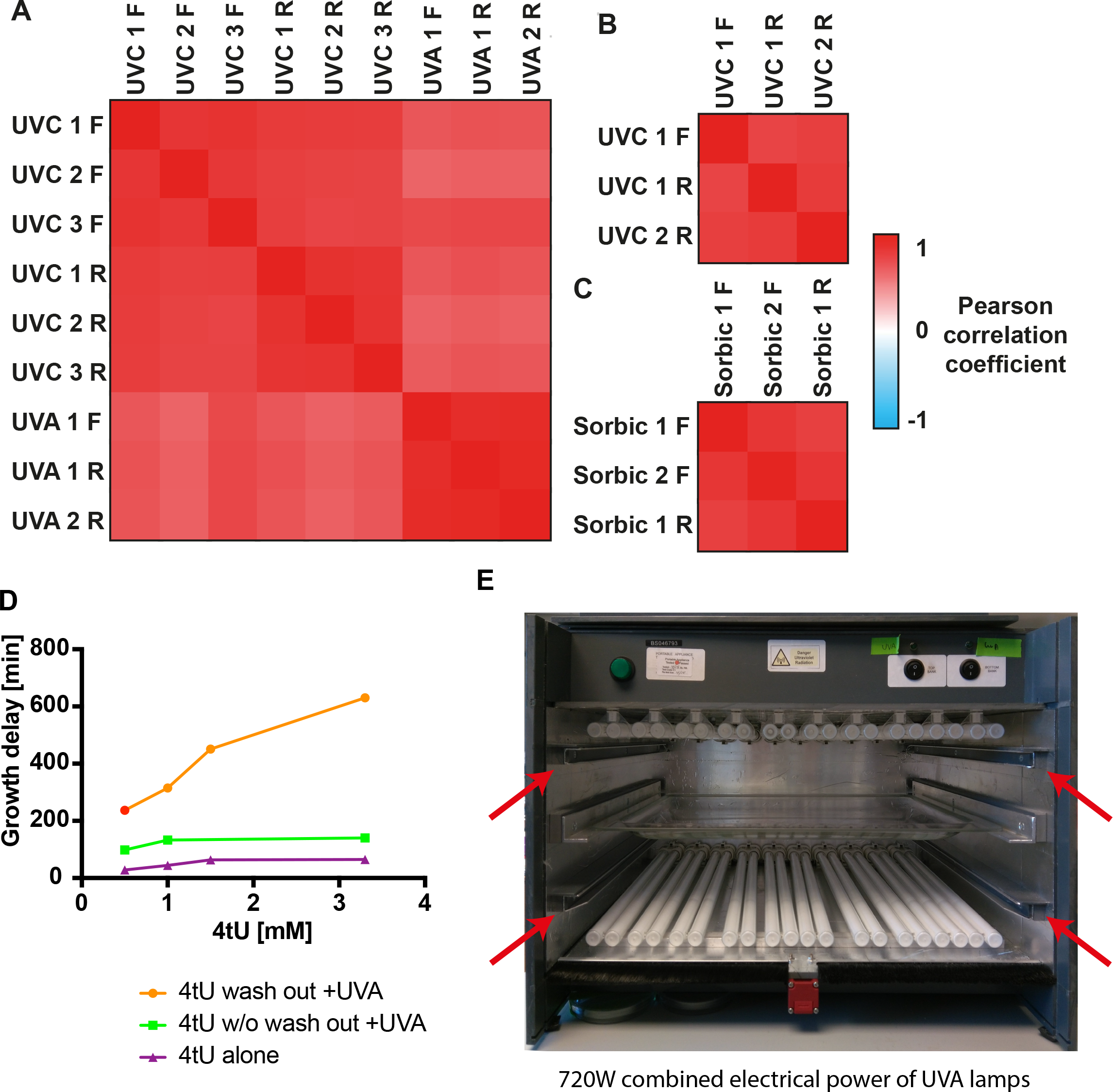
(A) Pairwise Pearson correlation coefficients for protein Log_2_ +UV/−UV ratios. *S. cerevisiae* 3 forward and 3 reverse isotopic labeling TRAPP repeats analyzed together with 1 forward and 2 reverse PAR-TRAPP repeats. (B) Analysis of *E. coli* 1 forward and 2 reverse TRAPP repeats performed as in (A). (C) Analysis of *S. cerevisiae* 1 forward and 2 reverse PAR-TRAPP experiments upon sorbic acid exposure performed as in (A). (D) Effect of 4tU and UVA treatments on the growth of yeast cells. Exponentially growing yeast cells were treated for 2 hours with 4-thiouracil at the indicated concentrations. The cultures were then irradiated with 350nm UVA light in the eBox for 30sec delivering 5.8J cm^-2^. The time taken by cultures to resume exponential growth was measured using tecan sunrise instrument. Samples: "4tU wash out +UVA" – growth delay of 4-thiouracil treated UVA irradiated cells, compared to UVA exposed sample. 4tU was removed prior to irradiation; " 4tU w/o wash out +UVA" – As sample 1, but 4tU persisted in the media while cells were irradiated; "4tU alone" – growth delay of cells treated with 4tU for 2h as compared to untreated cells without irradiation. (E) Frontal view on the eBox irradiation apparatus. The frontal door and the shutters are not present on the picture. Red arrows indicate rails for shutters, designed to prevent sample exposure to UV light while the lamps are warming up for stable UVA output. The UVA transparent sample tray made of borosilicate glass is placed between the two UVA lamp banks.

**Figure S2.**
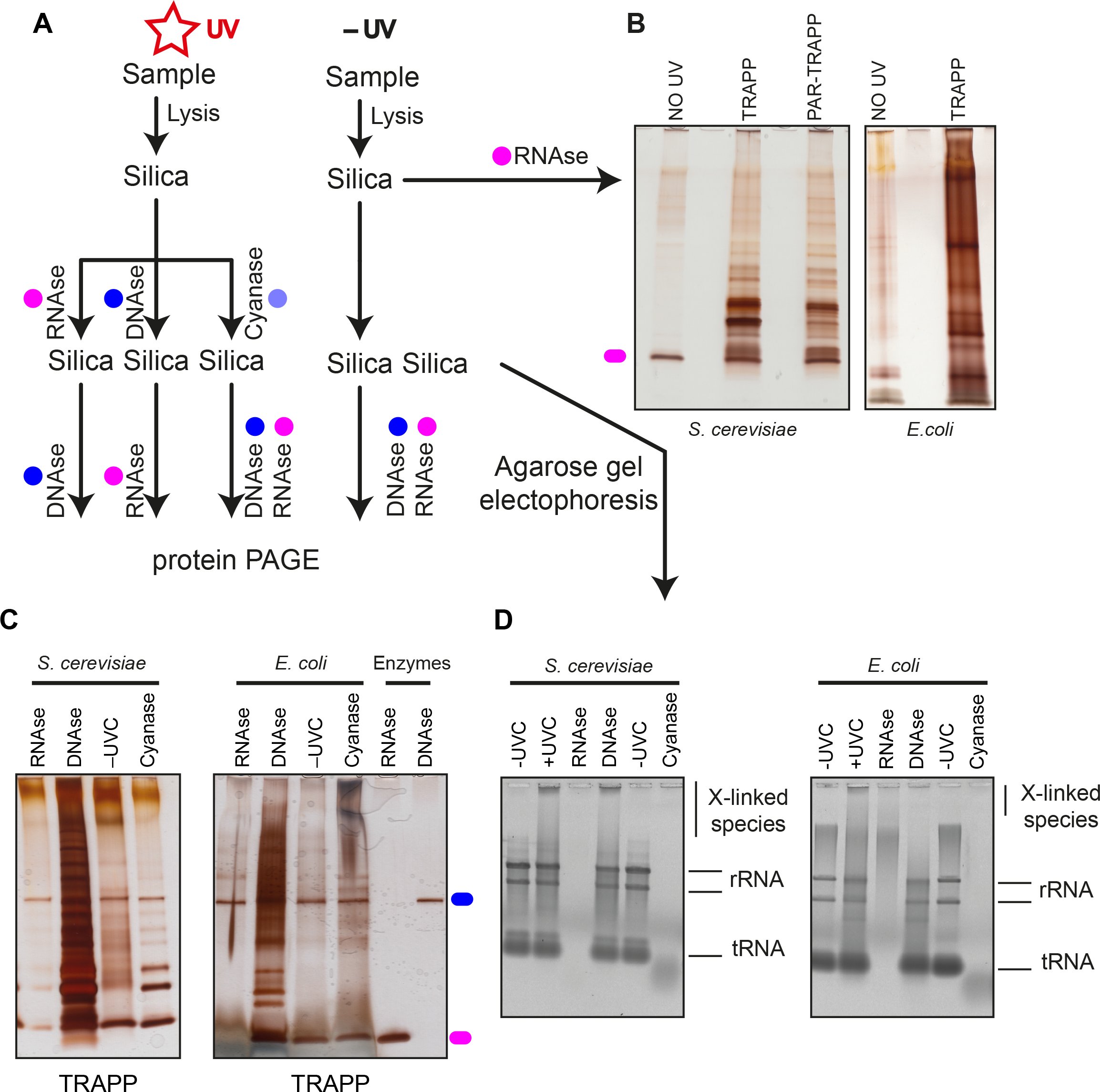
TRAPP protocol predominately recovers RNA-bound proteins. (A) The experimental setup indicating the stages when samples are collected. Colored circles designate treatment with the indicated enzyme. (B) Sample are purified following the TRAPP protocol as described in materials and methods. After RNase A and RNase T1 treatment to degrade the co-purifying RNA, sample was resolved on polyacrylamide gel and silver staining was performed. (C) TRAPP purified samples were treated with the indicated enzymes and loaded onto silica once again. After elution nucleic acids were resolved with agarose gel electrophoresis (see figure S2D), while the remainder of the sample was treated with the indicated enzyme followed by polyacrylamide gel electrophoresis and silver staining. (D) Same as in C, but the samples were collected before the second nuclease treatment and were then resolved on a SYBR Safe stained agarose gel.

**Figure S3.**
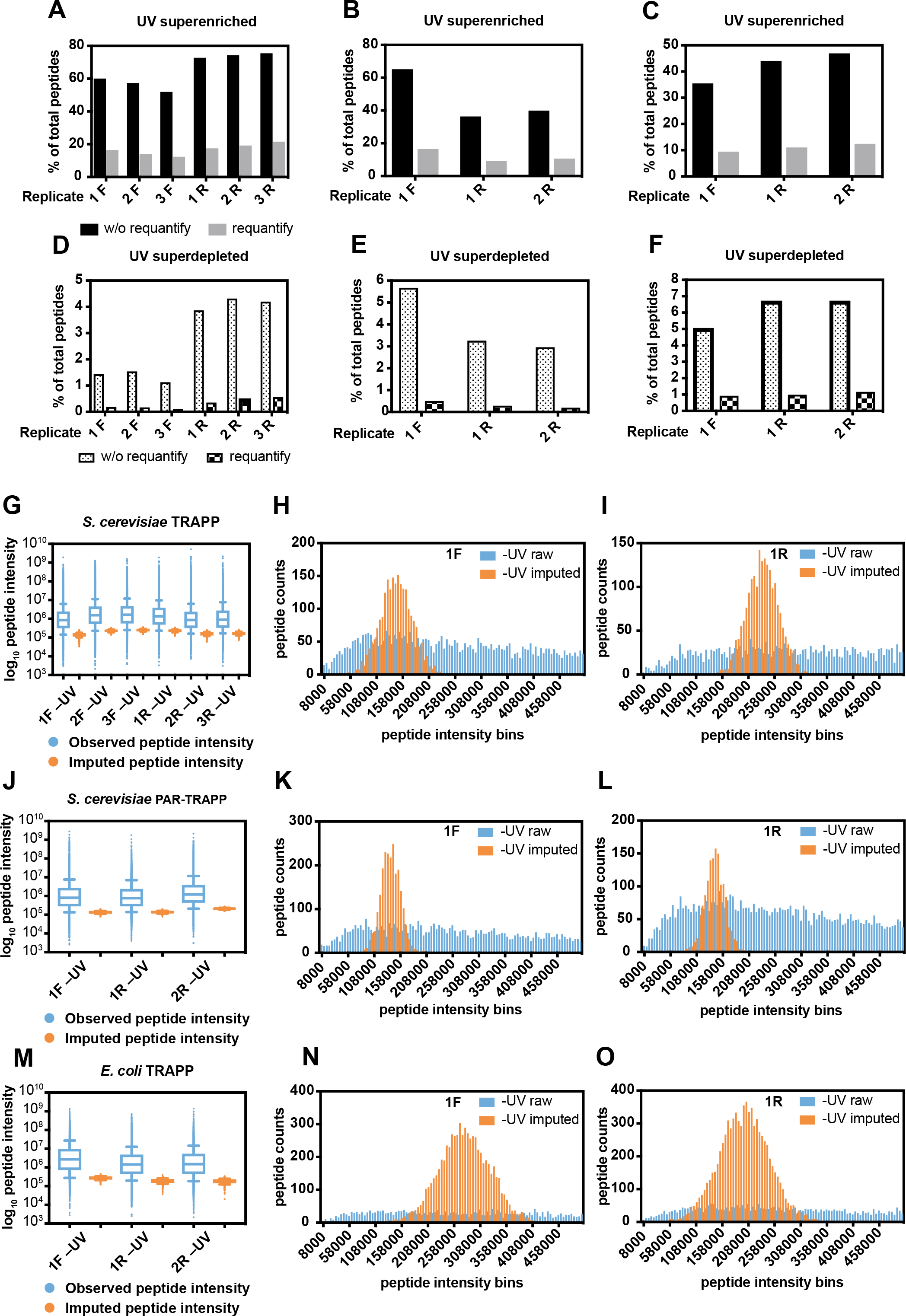
(A) The percentage of peptides with reported intensity in +UV sample, but not in −UV sample (superenriched peptides) by Maxquant in *S. cerevisiae* TRAPP SILAC quantification experiments without (black bars) or with (grey bars) "requantify" option enabled. 3 biological repeats had light-labelled cells UV irradiated (1F, 2F, 3F), while 3 other repeats (1R, 2R, 3R) had heavy-labelled cells UV irradiated. (B) The data of *S. cerevisiae* PAR-TRAPP experiments was analyzed the same way as in (A). (C) The data of *E. coli* TRAPP experiments was analyzed the same way as in (A), except the 2 biological repeats, which had light-labelled cells UV irradiated, were labelled 1R and 2R. (D) The percentage of peptides with reported intensity in – UV sample, but not in +UV sample (superdepleted peptides) by Maxquant in *S. cerevisiae* TRAPP without (dotted) or with (chequered) "requantify" option enabled. Sample labelling as in (A). (E) The data of *S. cerevisiae* PAR-TRAPP experiments was analyzed the same way as in (D). (F) The data of *E. coli* TRAPP experiments was analyzed the same way as in (D), sample labelling was as in figure (C). (G) Box plot of Log_10_ peptide intensity of −UV peptides from *S. cerevisiae* TRAPP (blue) samples (labelling as in (A)), plotted together with Log_10_ peptide intensity values imputed by imputeLCMD R package for −UV samples (orange). Box represent values between 25^th^ and 75^th^ percentiles, while whiskers represent 10^th^ and 90^th^ percentiles. All other data is represented as points below or above 10^th^ or 90^th^ percentiles respectively. Line inside the box represents median value. (H) Histogram of peptide intensity frequency obtained from −UV sample (1F), plotted for intensities from 0 to 5×10^5^ units. Color labelling is as in (G). (I) Same as figure (H), performed for sample 1R which had reversed SILAC labelling, compared to the sample analyzed in figure (H). (J) Box plot of Log_10_ peptide intensity of −UV peptides from *S. cerevisiae* PAR-TRAPP (blue) samples (labelling as in (B)), plotted together with Log_10_ peptide intensity values imputed by imputeLCMD R package for −UV samples (orange). (K) Histogram of peptide intensity frequency obtained from −UV sample (1F), plotted for intensities from 0 to 5×10^5^ units. Color labelling is as in (J). (L) Same analysis as figure (K), performed for sample 1R which had reversed SILAC labelling, compared to the sample analyzed in figure (K). (M) Box plot of Log_10_ peptide intensity of −UV peptides from *E. coli* TRAPP (blue) samples (labelling as in (C)), plotted together with Log_10_ peptide intensity values imputed by imputeLCMD R package for −UV samples (orange). (N) Histogram of peptide intensity frequency obtained from −UV sample (1F), plotted for intensities from 0 to 5×10^5^ units. Color labelling is as in (M). (O) Same analysis as figure (N), performed for sample 1R which had reversed SILAC labelling, compared to the sample analyzed in figure (N).

**Figure S4.**
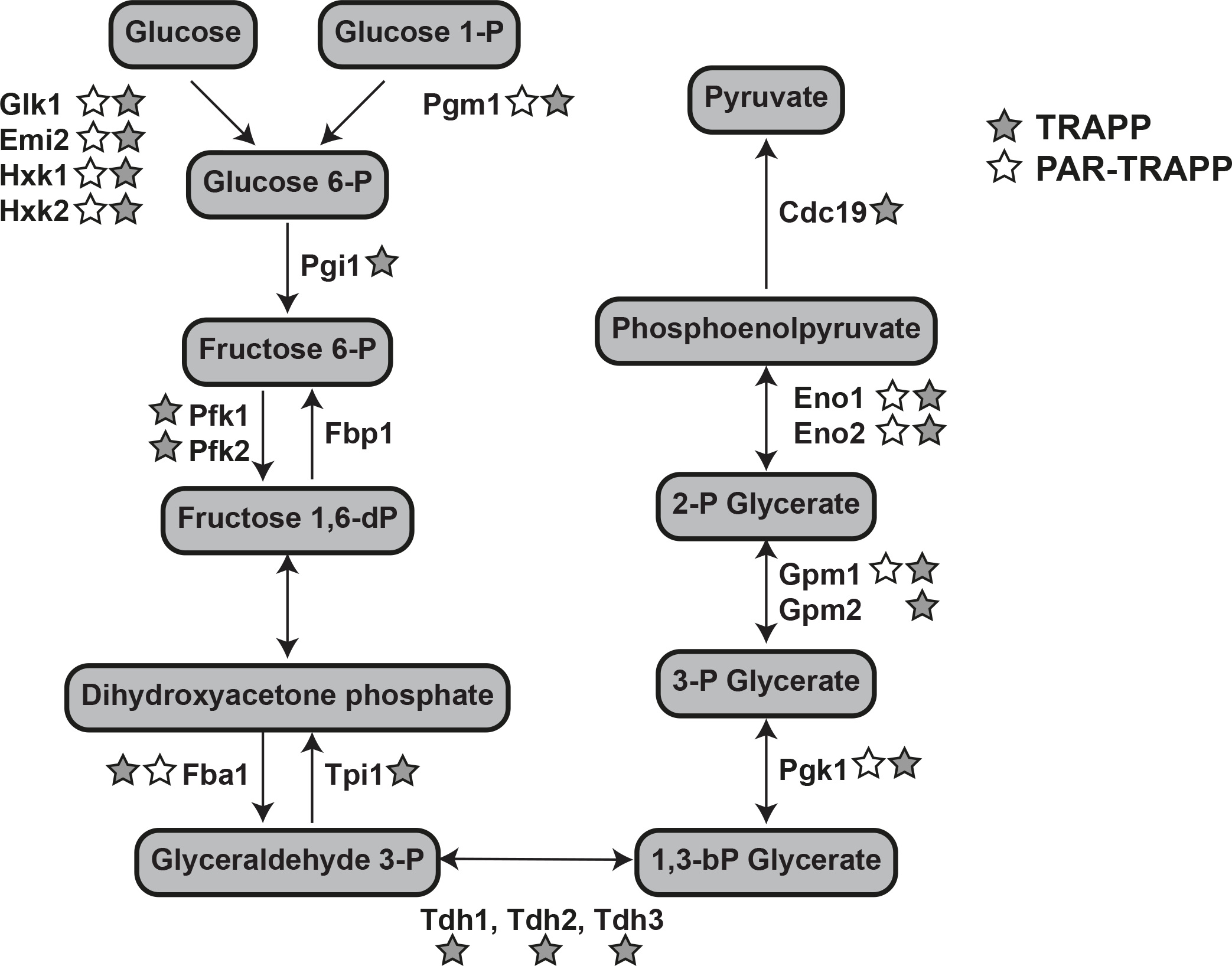
Glycolysis pathway in yeast *S. cerevisiae*, indicating intermediate metabolites and participating enzymes. Proteins identified as enriched by TRAPP and PAR-TRAPP are shown which grey and white stars respectively.

**Figure S5.**
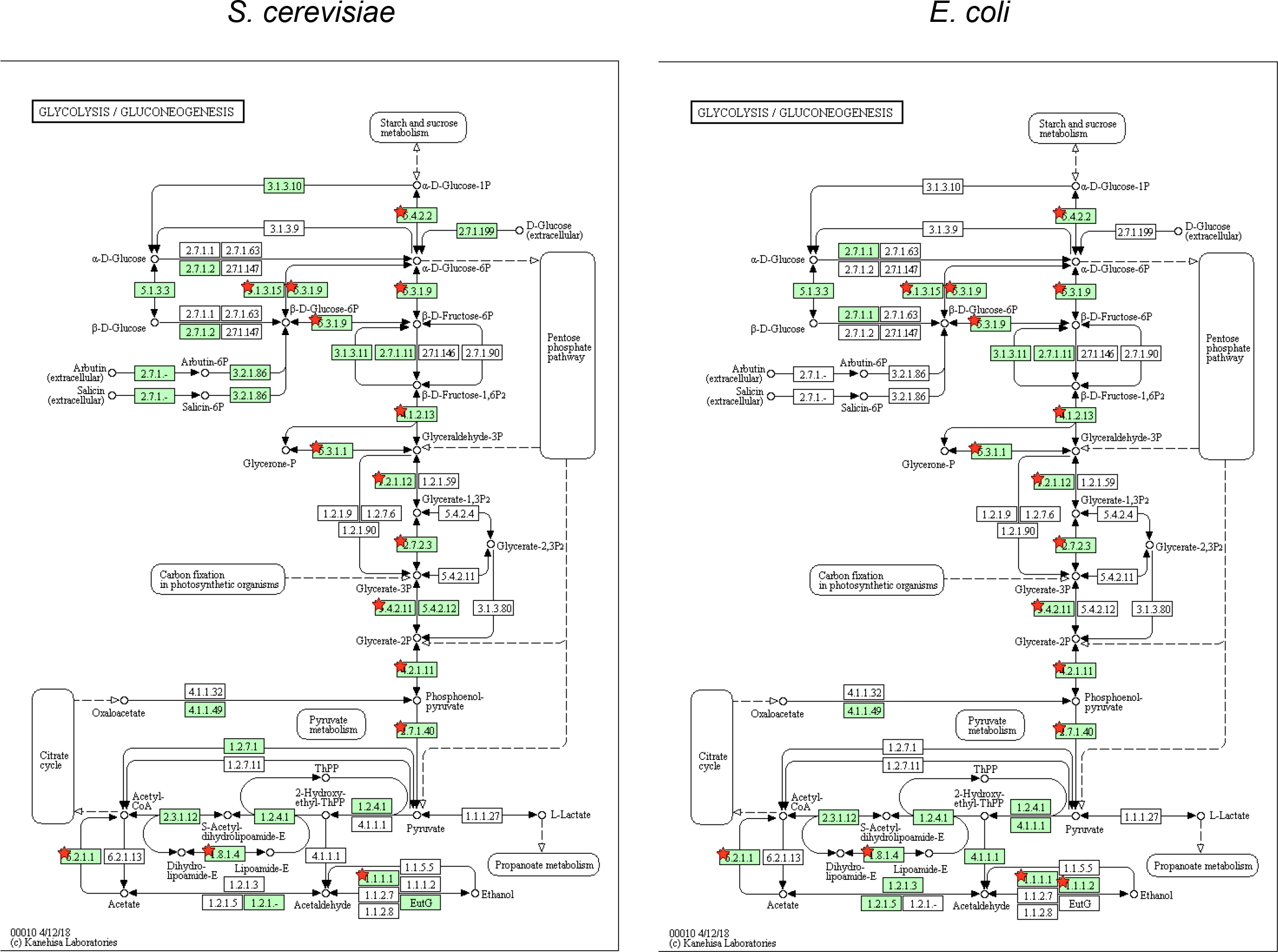
Schematic representation of Glycolysis/Gluconeogenesis KEGG pathway identified as enriched amongst conserved RNA interacting proteins of intermediary metabolism from yeast and bacteria (Figure 3C). Proteins, present in the indicated model organism are labelled in green. Red stars indicate proteins, identified as significantly enriched in TRAPP analysis.

**Figure S6.**
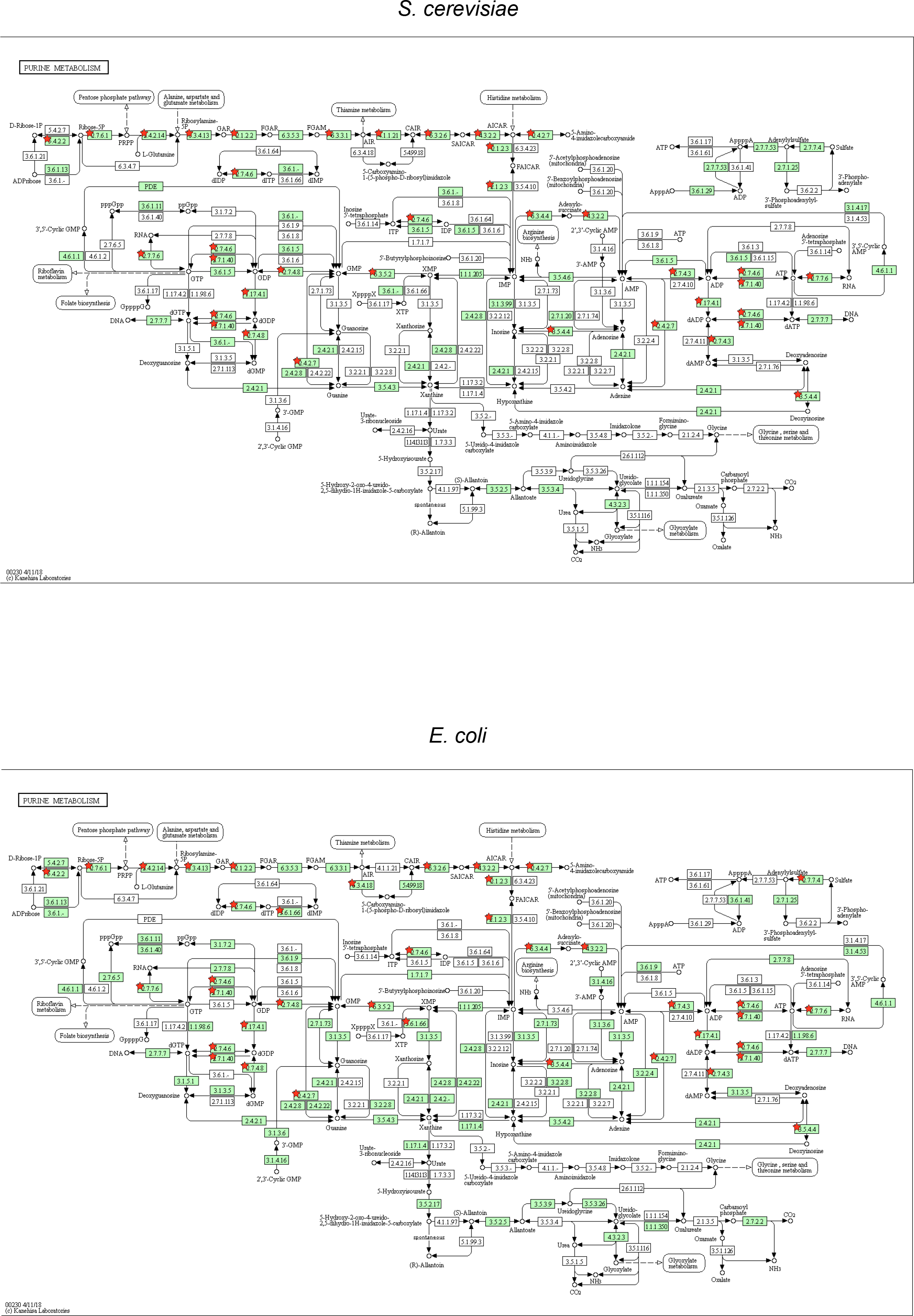
Schematic representation "Purine metabolism" KEGG pathway identified as enriched amongst conserved RNA interacting proteins of intermediary metabolism from yeast and bacteria (Figure 4C). Labelling as in figure S5.

**Figure S7.**
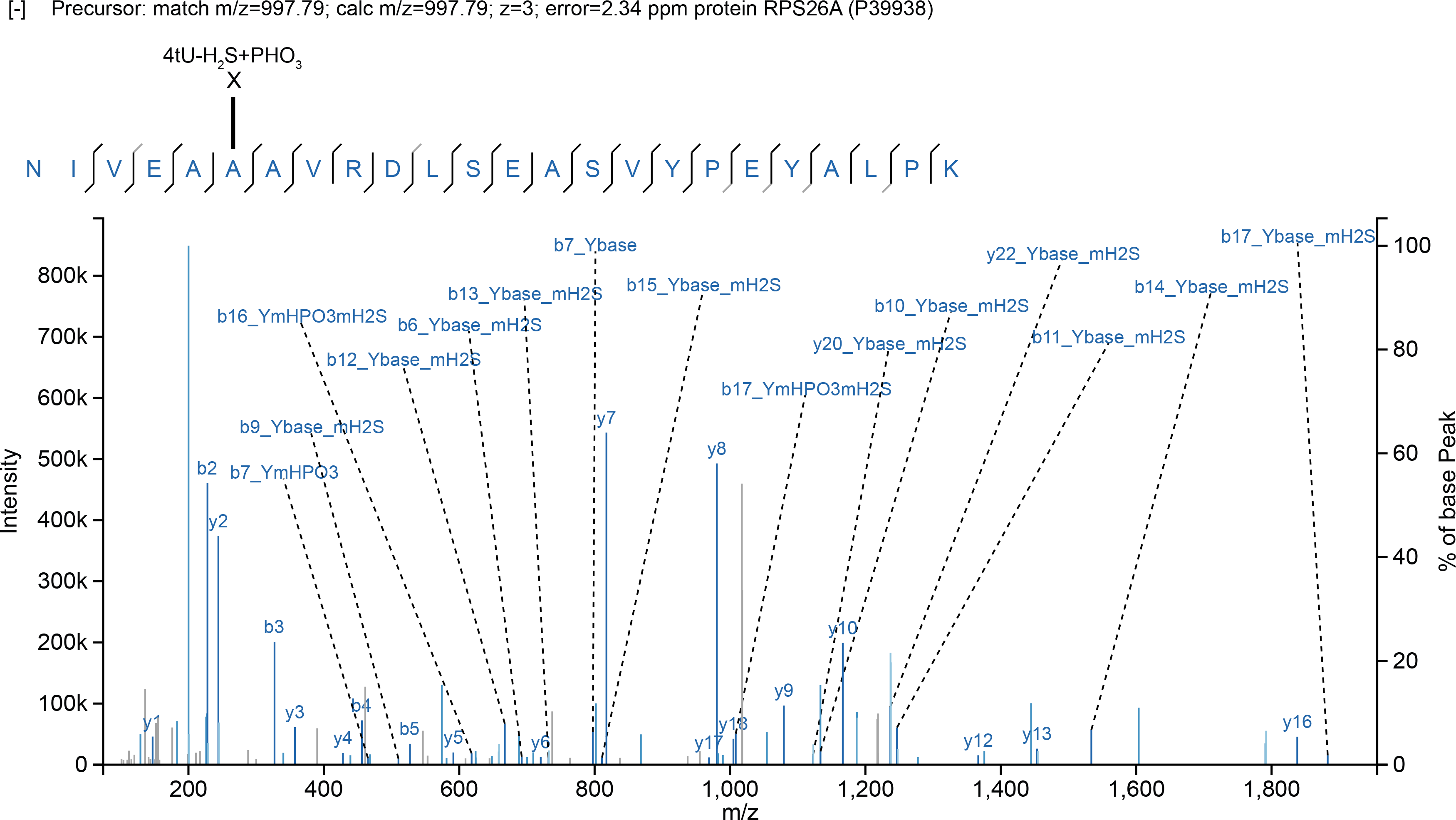
MS and MS/MS spectrum of peptide, crosslinked to 4-thiouridine via alanine residue. The observed peptide fragments are indicated on the peptide sequence with black bars. Grey bars indicate peptide fragments observed with an additional neutral loss. The software assigned site of crosslink is indicated above the peptide sequence. Allowed RNA fragments: Ybase - 4-thiouracil base; Ybase_mH2S - 4-thiouracil base with an H2S mass loss; YmHPO3mH2S - 4-thiouridine monophosphate with HPO3 and H2S mass losses; YmH2S - 4-thiouridine monophosphate with H2S mass loss; Y - 4-thiouridine monophosphate; YmHPO3 - 4-thiouridine monophosphate with HPO3 mass loss; YpHPO3mH2S - 4-thiouridine diphosphate with an H2S mass loss; YpHPO3 - 4-thiouridine diphosphate.

**Figure S8.**
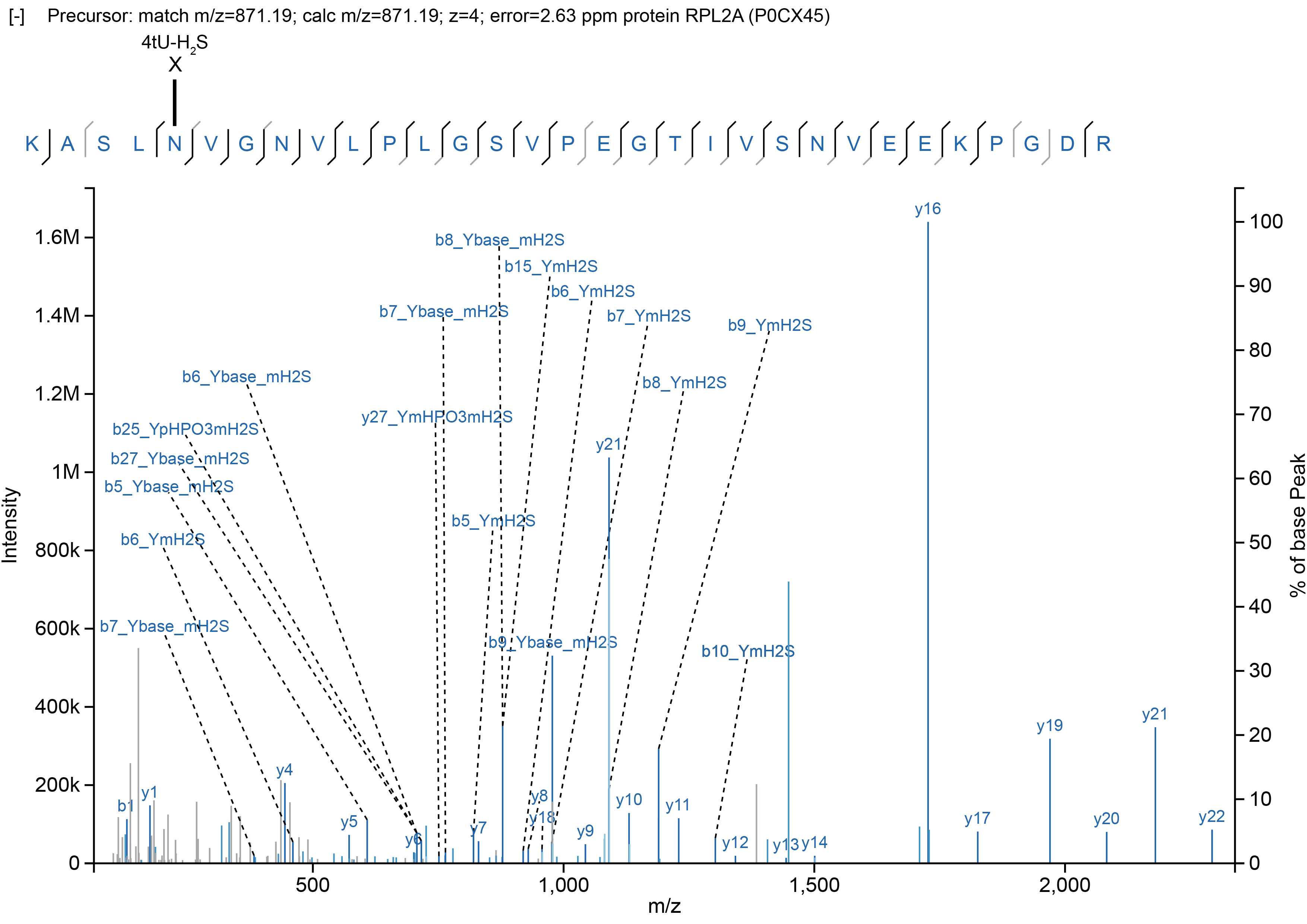
MS and MS/MS spectrum of peptide, crosslinked to 4-thiouridine via asparagine residue. The observed peptide fragments are indicated on the peptide sequence with black bars. Grey bars indicate peptide fragments observed with an additional neutral loss. The software assigned site of crosslink is indicated above the peptide sequence. Allowed RNA fragments: Ybase - 4-thiouracil base; Ybase_mH2S - 4-thiouracil base with an H2S mass loss; YmHPO3mH2S - 4-thiouridine monophosphate with HPO3 and H2S mass losses; YmH2S - 4-thiouridine monophosphate with H2S mass loss; Y - 4-thiouridine monophosphate; YmHPO3 - 4-thiouridine monophosphate with HPO3 mass loss; YpHPO3mH2S - 4-thiouridine diphosphate with an H2S mass loss; YpHPO3 - 4-thiouridine diphosphate. “P_” plus the RNA fragment denotes unfragmented peptide conjugated to an RNA fragment.

**Figure S9.**
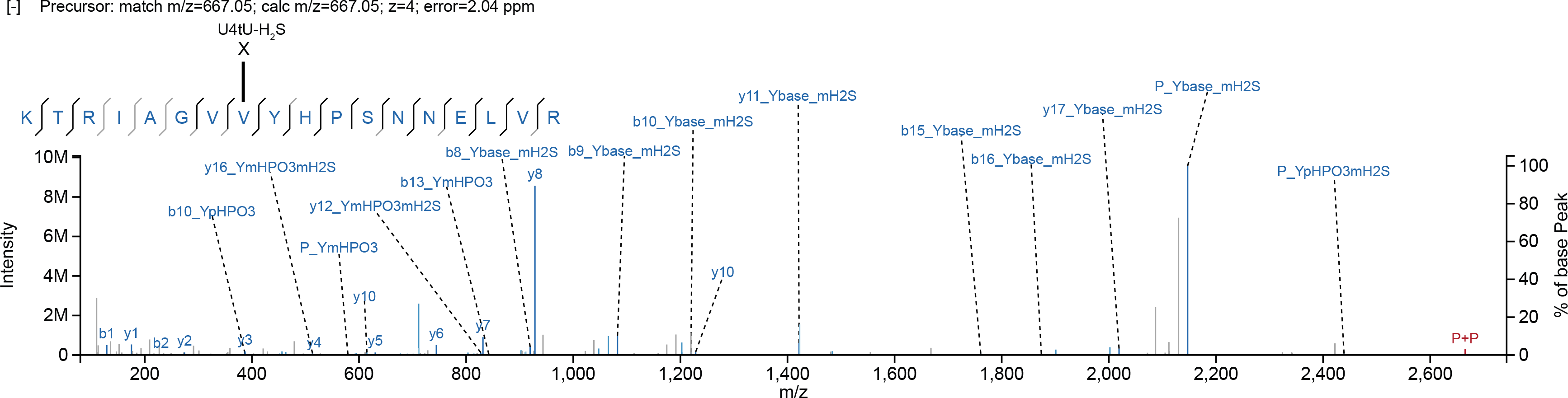
MS and MS/MS spectrum of peptide, crosslinked to 4-thiouridine via valine residue. The observed peptide fragments are indicated on the peptide sequence with black bars. Grey bars indicate peptide fragments observed with an additional neutral loss. The software assigned site of crosslink is indicated above the peptide sequence. Allowed RNA fragments: Ybase - 4-thiouracil base; Ybase_mH2S - 4-thiouracil base with an H2S mass loss; YmHPO3mH2S - 4-thiouridine monophosphate with HPO3 and H2S mass losses; YmH2S - 4-thiouridine monophosphate with H2S mass loss; Y - 4-thiouridine monophosphate; YmHPO3 - 4-thiouridine monophosphate with HPO3 mass loss; YpHPO3mH2S - 4-thiouridine diphosphate with an H2S mass loss; YpHPO3 - 4-thiouridine diphosphate. “P_” plus the RNA fragment denotes unfragmented peptide conjugated to an RNA fragment. “P+P” labels the unfragmented ion of Peptide conjugated to RNA. See also Supplementary Text.

**Table 1**: Numbers of RNA-protein crosslinks mapped using Xi

**Table S1:** List of the yeast *S. cerevisiae* proteins quantified with TRAPP

**Table S2**: List of the yeast *S. cerevisiae* proteins quantified with PAR-TRAPP

**Table S3**: List of Interpro protein domains identified as enriched in TRAPP data

**Table S4**: List of the yeast *E. coli* proteins quantified with TRAPP

**Table S5**: GO term analysis of RNA interacting proteins conserved between *S. cerevisiae* and E. coli

**Table S6**: List of *S. cerevisiae* proteins, quantified with PAR-TRAPP upon Sorbic acid exposure

**Table S7**: RNA species and their masses used in Xi targeted search for RNA-crosslinked peptides

**Table S8**: Peptides identified by the Xi search engine

**Table S9**: The analysis of the yeast proteins identified as crosslinked to RNA by Xi or RNPxl search engines, but not in PAR-TRAPP

**Table S10**: Crosslinking of Eno1 to mRNAs in CRAC analyses

## Supplementary Text

### Supplementary Text related to Figure 5

We would anticipate that the presence of a phosphate group would reduce RNA interactions due to its charge. However, it remained possible that the TiO_2_ enrichment might favour peptides that are simultaneously crosslinked and phosphorylated, increasing the apparent correlation. The precursor ion mass of a peptide crosslinked to 4-thiouridine 3′,5′-diphosphate is identical to the same peptide crosslinked to 4-thiouridine monophosphate but harboring a phospho-serine or threonine. If insufficient MS2 fragments coverage is obtained for such peptides, it becomes difficult to discriminate between the two possibilities. Excluding all crosslinks with 4-thiouridine diphosphate (4tU-H_2_S+HPO_3_), still revealed significant correlation between iTRAPP sites and reported protein phosphorylation. The P value was higher (1.3×10^-4^), reflecting the reduced number of crosslinks assessed.

### Comparison of RNP^xl^ and Xi

Comparison between RNPxl and Xi search engines on the mapping of RNA-protein crosslink spectra. In order to compare the performance of Xi with previously published software tools, we analyzed our data with the RNPxl package for proteome discoverer (Veit, Sachsenberg et al., 2016). The list of possible 4-thioU fragments and other search parameters were harmonized between Xi and RNPxl as much as possible and both search engines were used to analyze the same dataset (see Materials and Methods). Based on the FDR cut-off of 1% (see materials and methods) RNPxl was able to identify 1038 crosslink-containing peptide-spectrum matches (PSMs) of which 440 belonged to unique peptide–RNA crosslinks in 236 peptides from 120 protein groups (see Table S11). Thus, in our hands Xi performed better than RNPxl. However, we since RNPxl does not report hit to decoy database, we had to include the decoy sequences in the search database for RNPxl, which may have negatively affected the performance.

### Supplementary Text related to Figure S9

We noted that the iTRAPP data frequently contained ions mapped to unfragmented peptide, conjugated to fragmented RNA, illustrating the problem of RNA fragmentation at the expense of peptide fragmentation. The figure also illustrates the limitations of the current software, as it contains an ion annotated as unfragmented peptide, conjugated to 4-thiouridine lacking a phosphate group. Moreover, while a loss of phosphate is commonly observed, we speculate that it is unlikely to have a fragment conjugated to 4-thiouridine, while the precursor ion all the other fragments bear a 4-thioutidine with an H_2_S mass loss. Thus, it would be beneficial to include in the future releases a way to specify the list of possible RNA fragments based on the conjugated nucleotide, especially so for UVC crosslinking.

## References

Angelov, D., Stefanovsky, V., Dimitrov, S.I., Russanova, V.R., Keskinova, E., and Pashev, I.G. (1988). Protein-DNA crosslinking in reconstituted nucleohistone, nuclei and whole cells by picosecond UV laser irradiation. Nucleic Acids Res 16, 4525–4538.

Baltz, A.G., Munschauer, M., Schwanhausser, B., Vasile, A., Murakawa, Y., Schueler, M., Youngs, N., Penfold-Brown, D., Drew, K., Milek, M., et al. (2012). The mRNA-bound proteome and its global occupancy profile on protein-coding transcripts. Mol Cell 46, 674–690.

Bao, X., Guo, X., Yin, M., Tariq, M., Lai, Y., Kanwal, S., Zhou, J., Li, N., Lv, Y., Pulido-Quetglas, C., et al. (2018). Capturing the interactome of newly transcribed RNA. Nature Meth 15, 213.

Beckmann, B.M., Horos, R., Fischer, B., Castello, A., Eichelbaum, K., Alleaume, A.-M., Schwarzl, T., Curk, T., Foehr, S., Huber, W., et al. (2015). The RNA-binding proteomes from yeast to man harbour conserved enigmRBPs. Nature Comm 6, 10127.

Beckstead, A.A., Zhang, Y., de Vries, M.S., and Kohler, B. (2016). Life in the light: nucleic acid photoproperties as a legacy of chemical evolution. Phys Chem Chem Phys 18, 24228–24238.

Bley, C.J., Qi, X., Rand, D.P., Borges, C.R., Nelson, R.W., and Chen, J.J. (2011). RNA-protein binding interface in the telomerase ribonucleoprotein. Proc Natl Acad Sci USA 108, 20333–20338.

Boeynaems, S., Alberti, S., Fawzi, N.L., Mittag, T., Polymenidou, M., Rousseau, F., Schymkowitz, J., Shorter, J., Wolozin, B., Van Den Bosch, L., et al. (2018). Protein Phase Separation: A New Phase in Cell Biology. Trends in Cell Biology 28, 420–435.

Brescia, C.C., Kaw, M.K., and Sledjeski, D.D. (2004). The DNA binding protein H-NS binds to and alters the stability of RNA in vitro and in vivo. J Mol Biol 339, 505–514.

Buchan, J.R., Nissan, T., and Parker, R. (2010). Analyzing P-Bodies and Stress Granules in Saccharomyces cerevisiae. In Meth Enz (Academic Press), pp. 619–640.

Carpousis, A.J. (2007). The RNA degradosome of Escherichia coli: an mRNA-degrading machine assembled on RNase E. Ann Rev Microbiol 61, 71–87.

Cary, G.A., Vinh, D.B.N., May, P., Kuestner, R., and Dudley, A.M. (2015). Proteomic Analysis of Dhh1 Complexes Reveals a Role for Hsp40 Chaperone Ydj1 in Yeast P-Body Assembly. G3 5, 2497.

Castello, A., Fischer, B., Eichelbaum, K., Horos, R., Beckmann, B.M., Strein, C., Davey, N.E., Humphreys, D.T., Preiss, T., Steinmetz, L.M., et al. (2012). Insights into RNA biology from an atlas of mammalian mRNA-binding proteins. Cell 149, 1393–1406.

Castello, A., Fischer, B., Frese, Christian K., Horos, R., Alleaume, A.-M., Foehr, S., Curk, T., Krijgsveld, J., and Hentze, Matthias W. (2016). Comprehensive Identification of RNA-Binding Domains in Human Cells. Molecular Cell 63, 696–710.

Clemson, C.M., Hutchinson, J.N., Sara, S.A., Ensminger, A.W., Fox, A.H., Chess, A., and Lawrence, J.B. (2009). An architectural role for a nuclear noncoding RNA: NEAT1 RNA is essential for the structure of paraspeckles. Mol Cell 33, 717–726.

Clery, A., Sinha, R., Anczukow, O., Corrionero, A., Moursy, A., Daubner, G.M., Valcarcel, J., Krainer, A.R., and Allain, F.H. (2013). Isolated pseudo-RNA-recognition motifs of SR proteins can regulate splicing using a noncanonical mode of RNA recognition. Proc Natl Acad Sci USA 110, E2802–2811.

Cox, J., and Mann, M. (2008). MaxQuant enables high peptide identification rates, individualized p.p.b.-range mass accuracies and proteome-wide protein quantification. Nat Biotechnol 26, 1367–1372.

Cox, J., Neuhauser, N., Michalski, A., Scheltema, R.A., Olsen, J.V., and Mann, M. (2011). Andromeda: A Peptide Search Engine Integrated into the MaxQuant Environment. Journal of proteome research 10, 1794–1805.

Decker, C.J., and Parker, R. (2012). P-Bodies and Stress Granules: Possible Roles in the Control of Translation and mRNA Degradation. Cold Spring Harbor Perspectives in Biology 4.

Dodt, M., Roehr, J.T., Ahmed, R., and Dieterich, C. (2012). FLEXBAR—Flexible Barcode and Adapter Processing for Next-Generation Sequencing Platforms. Biology 1, 895–905.

Doneanu, C.E., Gafken, P.R., Bennett, S.E., and Barofsky, D.F. (2004). Mass spectrometry of UV-cross-linked protein-nucleic acid complexes: identification of amino acid residues in the single-stranded DNA-binding domain of human replication protein A. Anal Chem 76, 5667–5676.

Entelis, N., Brandina, I., Kamenski, P., Krasheninnikov, I.A., Martin, R.P., and Tarassov, I. (2006). A glycolytic enzyme, enolase, is recruited as a cofactor of tRNA targeting toward mitochondria in Saccharomyces cerevisiae. Genes & Development 20, 1609–1620.

Garcia-Moreno, M., Noerenberg, M., Ni, S., Jarvelin, A.I., Gonzalez-Almela, E., Lenz, C., Bach-Pages, M., Cox, V., Avolio, R., Davis, T., et al. (2018). Understanding RNP remodelling uncovers RBPs functionally required for viral replication. bioRxiv.

Gatto, L., Breckels, L.M., and Lilley, K.S. (2018). Assessing sub-cellular resolution in spatial proteomics experiments. bioRxiv.

Giese, S.H., Fischer, L., and Rappsilber, J. (2015). A study into the CID behavior of cross-linked peptides. Mol Cell Proteomics 15, 1094–1104.

Gilbert, C., Kristjuhan, A., Winkler, G.S., and Svejstrup, J.Q. (2004). Elongator interactions with nascent mRNA revealed by RNA immunoprecipitation. Mol Cell 14, 457–464.

Gilbert, C., and Svejstrup, J.Q. (2006). RNA immunoprecipitation for determining RNA-protein associations in vivo. Current protocols in molecular biology / edited by Frederick M Ausubel [et al.] Chapter 27, Unit 27 24.

Gomar-Alba, M., Jiménez-Martí, E., and del Olmo, M. (2012). The Saccharomyces cerevisiae Hot1p regulated gene YHR087W (HGI1) has a role in translation upon high glucose concentration stress. BMC Molecular Biology 13, 19.

Granneman, S., Kudla, G., Petfalski, E., and Tollervey, D. (2009). Identification of protein binding sites on U3 snoRNA and pre-rRNA by UV cross-linking and high-throughput analysis of cDNAs. Proceedings of the National Academy of Sciences of the United States of America 106, 9613–9618.

Granneman, S., Petfalski, E., Swiatkowska, A., and Tollervey, D. (2010). Cracking pre-40S ribosomal subunit structure by systematic analyses of RNA-protein cross-linking. The EMBO journal 29, 2026–2036.

Granneman, S., Petfalski, E., and Tollervey, D. (2011). A cluster of ribosome synthesis factors regulate pre-rRNA folding and 5.8S rRNA maturation by the Rat1 exonuclease. Embo J 30, 4006–4019.

Grosshans, H., Hurt, E., and Simos, G. (2000). An aminoacylation-dependent nuclear tRNA export pathway in yeast. Genes Dev 14, 830–840.

Harel-Sharvit, L., Eldad, N., Haimovich, G., Barkai, O., Duek, L., and Choder, M. (2010). RNA Polymerase II Subunits Link Transcription and mRNA Decay to Translation. Cell 143, 552–563.

Hasunuma, T., Sakamoto, T., and Kondo, A. (2016). Inverse metabolic engineering based on transient acclimation of yeast improves acid-containing xylose fermentation and tolerance to formic and acetic acids. Applied Microbiol Biotech 100, 1027–1038.

Hu, Z., Xia, B., Postnikoff, S.D.L., Shen, Z.-J., Tomoiaga, A.S., Harkness, T.A., Seol, J.H., Li, W., Chen, K., and Tyler, J.K. (2018). Ssd1 and Gcn2 suppress global translation efficiency in replicatively aged yeast while their activation extends lifespan. eLife 7, e35551.

Huang, B., Johansson, M.J., and Bystrom, A.S. (2005). An early step in wobble uridine tRNA modification requires the Elongator complex. RNA 11, 424–436.

Huang, R., Han, M., Meng, L., and Chen, X. (2018). Transcriptome-wide discovery of coding and noncoding RNA-binding proteins. Proc Natl Acad Sci USA 115, E3879.

Hulsen, T., de Vlieg, J., and Alkema, W. (2008). BioVenn - a web application for the comparison and visualization of biological lists using area-proportional Venn diagrams. BMC Genomics 9, 488.

Hurt, E., Luo, M.J., Rother, S., Reed, R., and Strasser, K. (2004). Cotranscriptional recruitment of the serine-arginine-rich (SR)-like proteins Gbp2 and Hrb1 to nascent mRNA via the TREX complex. Proc Natl Acad Sci USA 101, 1858–1862.

Jain, S., Wheeler, J.R., Walters, R.W., Agrawal, A., Barsic, A., and Parker, R. (2016). ATPase modulated stress granules contain a diverse proteome and substructure. Cell 164, 487–498.

Jiao, X., Sherman, B.T., Huang, D.W., Stephens, R., Baseler, M.W., Lane, H.C., and Lempicki, R.A. (2012). DAVID-WS: a stateful web service to facilitate gene/protein list analysis. Bioinformatics 28, 1805–1806.

Jin, M., Fuller, G.G., Han, T., Yao, Y., Alessi, A.F., Freeberg, M.A., Roach, N.P., Moresco, J.J., Karnovsky, A., Baba, M., et al. (2017). Glycolytic Enzymes Coalesce in G Bodies under Hypoxic Stress. Cell Rep 20, 895–908.

Kamada, Y. (2017). Novel tRNA function in amino acid sensing of yeast Tor complex1. Genes to Cells 22, 135–147.

Kappel, L., Loibl, M., Zisser, G., Klein, I., Fruhmann, G., Gruber, C., Unterweger, S., Rechberger, G., Pertschy, B., and Bergler, H. (2012). Rlp24 activates the AAA-ATPase Drg1 to initiate cytoplasmic pre-60S maturation. J Cell Biol 199, 771–782.

Kawahata, M., Masaki, K., Fujii, T., and Iefuji, H. (2006). Yeast genes involved in response to lactic acid and acetic acid: acidic conditions caused by the organic acids in Saccharomyces cerevisiae cultures induce expression of intracellular metal metabolism genes regulated by Aft1p. FEMS yeast research 6, 924–936.

Kramer, K., Hummel, P., Hsiao, H.-H., Luo, X., Wahl, M., and Urlaub, H. (2011). Mass-spectrometric analysis of proteins cross-linked to 4-thio-uracil- and 5-bromo-uracil-substituted RNA. Internat J Mass Spec 304, 184–194.

Kramer, K., Sachsenberg, T., Beckmann, B.M., Qamar, S., Boon, K.-L., Hentze, M.W., Kohlbacher, O., and Urlaub, H. (2014). Photo-cross-linking and high-resolution mass spectrometry for assignment of RNA-binding sites in RNA-binding proteins. Nat Meth 11, 1064–1070.

Kurischko, C., Kim, H.K., Kuravi, V.K., Pratzka, J., and Luca, F.C. (2011). The yeast Cbk1 kinase regulates mRNA localization via the mRNA-binding protein Ssd1. The Journal of Cell Biology 192, 583.

Kwon, S.C., Yi, H., Eichelbaum, K., Fohr, S., Fischer, B., You, K.T., Castello, A., Krijgsveld, J., Hentze, M.W., and Kim, V.N. (2013). The RNA-binding protein repertoire of embryonic stem cells. Nat Struct Mol Biol 20, 1122–1130.

Langdon, E.M., Qiu, Y., Ghanbari Niaki, A., McLaughlin, G.A., Weidmann, C.A., Gerbich, T.M., Smith, J.A., Crutchley, J.M., Termini, C.M., Weeks, K.M., et al. (2018). mRNA structure determines specificity of a polyQ-driven phase separation. Science 360, 922.

Lazar, C., Gatto, L., Ferro, M., Bruley, C., and Burger, T. (2016). Accounting for the Multiple Natures of Missing Values in Label-Free Quantitative Proteomics Data Sets to Compare Imputation Strategies. J Proteome Res 15, 1116–1125.

Lewis, J., and Tollervey, D. (2000). Like attracts like: Getting RNA processing together in the nucleus. Science 288, 1385–1389.

Lin, Y., Protter, David S.W., Rosen, Michael K., and Parker, R. (2015). Formation and Maturation of Phase-Separated Liquid Droplets by RNA-Binding Proteins. Molecular Cell 60, 208–219.

Lo, K.Y., Li, Z., Bussiere, C., Bresson, S., Marcotte, E.M., and Johnson, A.W. (2010). Defining the pathway of cytoplasmic maturation of the 60S ribosomal subunit. Mol Cell 39, 196–208.

Lotan, R., Goler-Baron, V., Duek, L., Haimovich, G., and Choder, M. (2007). The Rpb7p subunit of yeast RNA polymerase II plays roles in the two major cytoplasmic mRNA decay mechanisms. J Cell Biol 178, 1133–1143.

Macvanin, M., Edgar, R., Cui, F., Trostel, A., Zhurkin, V., and Adhya, S. (2012). Noncoding RNAs Binding to the Nucleoid Protein HU in Escherichia coli. J Bact 194, 6046–6055.

Maharana, S., Wang, J., Papadopoulos, D.K., Richter, D., Pozniakovsky, A., Poser, I., Bickle, M., Rizk, S., Guillén-Boixet, J., Franzmann, T., et al. (2018). RNA buffers the phase separation behavior of prion-like RNA binding proteins. Science 359, 918–921.

Maly, P., Rinke, J., Ulmer, E., Zwieb, C., and Brimacombe, R. (1980). Precise localization of the site of cross-linking between protein L4 and 23S ribonucleic acid induced by mild ultraviolet irradiation of Escherichia coli 50S ribosomal subunits. Biochemistry 19, 4179–4188.

Mendes, M.L., Fischer, L., Chen, Z.A., Barbon, M., Reilly, F.J., Bohlke-Schneider, M., Belsom, A., Dau, T., Combe, C.W., Graham, M., et al. (2018). An integrated workflow for cross-linking/mass spectrometry. bioRxiv.

Mira, N.P., Lourenço, A.B., Fernandes, A.R., Becker, J.D., and Sá-Correia, I. (2008). The RIM101 pathway has a role in Saccharomyces cerevisiae adaptive response and resistance to propionic acid and other weak acids. FEMS yeast research 9, 202–216.

Mital, R., Albrecht, U., and Schumperli, D. (1993). Detection of UV-induced RNA:protein crosslinks in snRNPs by oligonucleotides complementary to the snRNA. Nucleic Acids Res 21, 1049–1050.

Miura, N., Shinohara, M., Tatsukami, Y., Sato, Y., Morisaka, H., Kuroda, K., and Ueda, M. (2013). Spatial Reorganization of Saccharomyces cerevisiae Enolase To Alter Carbon Metabolism under Hypoxia. Eukaryotic cell 12, 1106–1119.

Motamedi, M.R., Verdel, A., Colmenares, S.U., Gerber, S.A., Gygi, S.P., and Moazed, D. (2004). Two RNAi complexes, RITS and RDRC, physically interact and localize to noncoding centromeric RNAs. Cell 119, 789–802.

Munder, M.C., Midtvedt, D., Franzmann, T., Nüske, E., Otto, O., Herbig, M., Ulbricht, E., Müller, P., Taubenberger, A., Maharana, S., et al. (2016). A pH-driven transition of the cytoplasm from a fluid- to a solid-like state promotes entry into dormancy. eLife 5, e09347.

Niranjanakumari, S., Lasda, E., Brazas, R., and Garcia-Blanco, M.A. (2002). Reversible cross-linking combined with immunoprecipitation to study RNA-protein interactions in vivo. Methods 26, 182–190.

Olsen, J.V., Macek, B., Lange, O., Makarov, A., Horning, S., and Mann, M. (2007). Higher-energy C-trap dissociation for peptide modification analysis. Nature Methods 4, 709.

Ong, S.E., Blagoev, B., Kratchmarova, I., Kristensen, D.B., Steen, H., Pandey, A., and Mann, M. (2002). Stable isotope labeling by amino acids in cell culture, SILAC, as a simple and accurate approach to expression proteomics. Mol Cell Proteomics 1, 376–386.

Pertschy, B., Saveanu, C., Zisser, G., Lebreton, A., Tengg, M., Jacquier, A., Liebminger, E., Nobis, B., Kappel, L., van der Klei, I., et al. (2007). Cytoplasmic Recycling of 60S Preribosomal Factors Depends on the AAA Protein Drg1. Mol Cell Biol 27, 6581–6592.

Ramirez-Gaona, M., Marcu, A., Pon, A., Guo, An C., Sajed, T., Wishart, N.A., Karu, N., Djoumbou Feunang, Y., Arndt, D., and Wishart, D.S. (2017). YMDB 2.0: a significantly expanded version of the yeast metabolome database. Nucleic Acids Research 45, D440–D445.

Rappsilber, J., Ishihama, Y., and Mann, M. (2003). Stop and go extraction tips for matrix-assisted laser desorption/ionization, nanoelectrospray, and LC/MS sample pretreatment in proteomics. Anal Chem 75, 663–670.

Reidegeld, K.A., Eisenacher, M., Kohl, M., Chamrad, D., Körting, G., Blüggel, M., Meyer, H.E., and Stephan, C. (2008). An easy-to-use Decoy Database Builder software tool, implementing different decoy strategies for false discovery rate calculation in automated MS/MS protein identifications. Proteomics 8, 1129–1137.

Rhode, B.M., Hartmuth, K., Urlaub, H., and Luhrmann, R. (2003). Analysis of site-specific protein-RNA cross-links in isolated RNP complexes, combining affinity selection and mass spectrometry. RNA 9, 1542–1551.

Riback, J.A., Katanski, C.D., Kear-Scott, J.L., Pilipenko, E.V., Rojek, A.E., Sosnick, T.R., and Drummond, D.A. (2017). Stress-triggered phase separation is an adaptive, evolutionarily tuned response. Cell 168, 1028–1040.e1019.

Richter, F.M., Hsiao, H.H., Plessmann, U., and Urlaub, H. (2009). Enrichment of protein-RNA crosslinks from crude UV-irradiated mixtures for MS analysis by on-line chromatography using titanium dioxide columns. Biopolymers 91, 297–309.

Ritchie, M.E., Phipson, B., Wu, D., Hu, Y., Law, C.W., Shi, W., and Smyth, G.K. (2015). limma powers differential expression analyses for RNA-sequencing and microarray studies. Nucleic Acids Res 43, e47–e47.

Sajed, T., Marcu, A., Ramirez, M., Pon, A., Guo, A.C., Knox, C., Wilson, M., Grant, J.R., Djoumbou, Y., and Wishart, D.S. (2016). ECMDB 2.0: A richer resource for understanding the biochemistry of E. coli. Nucleic Acids Research 44, D495–D501.

Sen, N.D., Zhou, F., Harris, M.S., Ingolia, N.T., and Hinnebusch, A.G. (2016). eIF4B stimulates translation of long mRNAs with structured 5′ UTRs and low closed-loop potential but weak dependence on eIF4G. Proc Natl Acad Sci USA.

Shetlar, M.D. (1980). Cross-Linking of Proteins to Nucleic Acids by Ultraviolet Light. In Photochemical and Photobiological Reviews, K.C. Smith, ed. (Springer US), pp. 105–197.

Shevchenko, A., Wilm, M., and Mann, M. (1997). Peptide sequencing by mass spectrometry for homology searches and cloning of genes. J Protein Chem 16, 481–490.

Sonnhammer, E.L.L., and Östlund, G. (2015). InParanoid 8: orthology analysis between 273 proteomes, mostly eukaryotic. Nucleic Acids Res 43, D234–D239.

The UniProt Consortium (2017). UniProt: the universal protein knowledgebase. Nucleic Acids Res 45, D158–D169.

Trendel, J., Schwarzl, T., Prakash, A., Bateman, A., Hentze, M.W., and Krijgsveld, J. (2018). The Human RNA-Binding Proteome and Its Dynamics During Arsenite-Induced Translational Arrest. bioRxiv.

Tuck, A.C., and Tollervey, D. (2013). A transcriptome-wide atlas of RNP composition reveals diverse classes of mRNAs and lncRNAs. Cell 154, 996–1009.

Tyanova, S., Temu, T., Sinitcyn, P., Carlson, A., Hein, M.Y., Geiger, T., Mann, M., and Cox, J. (2016). The Perseus computational platform for comprehensive analysis of (prote)omics data. Nature Methods 13, 731.

Urlaub, H., Hartmuth, K., Kostka, S., Grelle, G., and Luhrmann, R. (2000). A general approach for identification of RNA-protein cross-linking sites within native human spliceosomal small nuclear ribonucleoproteins (snRNPs). Analysis of RNA-protein contacts in native U1 and U4/U6.U5 snRNPs. J Biol Chem 275, 41458–41468.

van der Lee, R., Buljan, M., Lang, B., Weatheritt, R.J., Daughdrill, G.W., Dunker, A.K., Fuxreiter, M., Gough, J., Gsponer, J., Jones, D.T., et al. (2014). Classification of Intrinsically Disordered Regions and Proteins. Chemical Reviews 114, 6589–6631.

Van Nostrand, E.L., Pratt, G.A., Shishkin, A.A., Gelboin-Burkhart, C., Fang, M.Y., Sundararaman, B., Blue, S.M., Nguyen, T.B., Surka, C., Elkins, K., et al. (2016). Robust transcriptome-wide discovery of RNA binding protein binding sites with enhanced CLIP (eCLIP). Nature Meth 13, 508–514.

Veit, J., Sachsenberg, T., Chernev, A., Aicheler, F., Urlaub, H., and Kohlbacher, O. (2016). LFQProfiler and RNPxl: Open-Source Tools for Label-Free Quantification and Protein–RNA Cross-Linking Integrated into Proteome Discoverer. J Proteome Res 15, 3441–3448.

Vizcaíno, J.A., Csordas, A., del-Toro, N., Dianes, J.A., Griss, J., Lavidas, I., Mayer, G., Perez-Riverol, Y., Reisinger, F., Ternent, T., et al. (2016). 2016 update of the PRIDE database and its related tools. Nucleic Acids Research 44, D447–D456.

Wagenmakers, A.J., Reinders, R.J., and van Venrooij, W.J. (1980). Cross-linking of mRNA to proteins by irradiation of intact cells with ultraviolet light. European journal of biochemistry / FEBS 112, 323–330.

Wang, M., Herrmann, C.J., Simonovic, M., Szklarczyk, D., and Mering, C. (2015). Version 4.0 of PaxDb: Protein abundance data, integrated across model organisms, tissues, and cell-lines. Proteomics 15, 3163–3168.

Webb, S., Hector, R.D., Kudla, G., and Granneman, S. (2014). PAR-CLIP data indicate that Nrd1-Nab3-dependent transcription termination regulates expression of hundreds of protein coding genes in yeast. Genome Biol 15, R8.

Weigert, C., Steffler, F., Kurz, T., Shellhammer, T.H., and Methner, F.-J. (2009). Application of a Short Intracellular pH Method to Flow Cytometry for Determining Saccharomyces cerevisiae Vitality. Applied Env Microbiol 75, 5615–5620.

Windbichler, N., von Pelchrzim, F., Mayer, O., Csaszar, E., and Schroeder, R. (2008). Isolation of small RNA-binding proteins from E. coli: Evidence for frequent interaction of RNAs with RNA polymerase. RNA Biology 5, 30–40.

Wiśniewski, J.R., Zielinska, D.F., and Mann, M. (2011). Comparison of ultrafiltration units for proteomic and N-glycoproteomic analysis by the filter-aided sample preparation method. Analytical Biochemistry 410, 307–309.

Woolford, J.L., and Baserga, S.J. (2013). Ribosome Biogenesis in the Yeast Saccharomyces cerevisiae. Genetics 195, 643–681.

Wu, Z.A., Murphy, C., Callan, H.G., and Gall, J.G. (1991). Small nuclear ribonucleoproteins and heterogeneous nuclear ribonucleoproteins in the amphibian germinal vesicle: loops, spheres, and snurposomes. J Cell Biol 113, 465–483.

Yan, J.X., Wait, R., Berkelman, T., Harry, R.A., Westbrook, J.A., Wheeler, C.H., and Dunn, M.J. (2000). A modified silver staining protocol for visualization of proteins compatible with matrix-assisted laser desorption/ionization and electrospray ionization-mass spectrometry. Electrophoresis 21, 3666–3672.

Zhang, A., Derbyshire, V., Salvo, J.L., and Belfort, M. (1995). Escherichia coli protein StpA stimulates self-splicing by promoting RNA assembly in vitro. RNA 1, 783–793.

Zou, X., Dai, X., Liu, K., Zhao, H., Song, D., and Su, H. (2014). Photophysical and Photochemical Properties of 4-Thiouracil: Time-Resolved IR Spectroscopy and DFT Studies. J Phys Chem B 118, 5864–5872.

